# *Egr2* induction in *Drd1*^+^ ensembles of the ventrolateral striatum supports the development of cocaine reward

**DOI:** 10.1101/2020.11.26.400721

**Authors:** Diptendu Mukherjee, Ben Jerry Gonzales, Reut Ashwal-Fluss, Hagit Turm, Maya Groysman, Ami Citri

**Author notes:** Authors contributed equally (Diptendu Mukherjee, Ben Jerry Gonzales).

## Abstract

Drug addiction develops due to brain-wide plasticity within neuronal ensembles, mediated by dynamic gene expression. Though the most common approach to identify such ensembles relies on immediate early gene expression, little is known of how the activity of these genes is linked to modified behavior observed following repeated drug exposure. To address this gap, we present a broad-to-specific approach, beginning with a comprehensive investigation of brain-wide cocaine-driven gene expression, through the description of dynamic spatial patterns of gene induction in subregions of the striatum, and finally address functionality of region-specific gene induction in the development of cocaine preference. Our findings reveal differential cell-type specific dynamic transcriptional recruitment patterns within two subdomains of the dorsal striatum following repeated cocaine exposure. Furthermore, we demonstrate that induction of the IEG Egr2 in the ventrolateral striatum, as well as the cells within which it is expressed, are required for the development of cocaine seeking.

**Impact statement:** VLS ensembles are dynamically recruited by cocaine experiences to mediate cocaine reward.

## Introduction

Psychostimulant addiction is characterized by life-long behavioral abnormalities, driven by circuit-specific modulation of gene expression (Nestler, 2014; Nestler and Lüscher, 2019; Salery et al., 2020; Steiner, 2016). Induction of immediate-early gene (IEG) transcription in the nucleus accumbens (NAc) and dorsal striatum (DS) are hallmarks of psychostimulant exposure (Caprioli et al., 2017; Chandra and Lobo, 2017; Gao et al., 2017b; Gonzales et al., 2020; Guez-Barber et al., 2011; Hope et al., 1994; Mukherjee et al., 2018; Nestler et al., 1993; Nestler, 2001; Nestler and Aghajanian, 1997; Piechota et al., 2010; Turm et al., 2014). As such, IEG induction has been utilized to support the identification of functional neuronal assemblies mediating the development of cocaine-elicited behaviors [“cocaine ensembles”; (Bobadilla et al., 2020; Cruz et al., 2013)]. Within these striatal structures, the principal neuronal type is the spiny projection neuron (SPN), which is comprised of two competing subtypes, defined by their differential expression of dopamine receptors. Expression of the D1R dopamine receptor is found on direct-pathway neurons, responsible for action selection by promoting behavioral responses, while D2R-expressing indirect pathway neurons are responsible for action selection through behavioral inhibition (Kreitzer and Malenka, 2008; Lipton et al., 2019). With regard to the cellular composition of cocaine ensembles, in the NAc, Fos-expressing cocaine ensembles have been found to be enriched for D1R expression (Koya et al., 2009). Within the DS, IEG expression is spatially segregated to the medial (MS) and ventrolateral (VLS) subdomains (Gonzales et al., 2020; Steiner and Gerfen, 1993). While psychostimulant-responsive ensembles in the MS have been found to encompass both D1R^+^ and D2R^+^ neurons (Caprioli et al., 2017; Cruz et al., 2015; Gonzales et al., 2020; Li et al., 2015; Rubio et al., 2015), in the VLS, these are enriched for D1R expression (Steiner 1993, Gonzales 2020).

Depending on the history of prior cocaine exposure, a unique pattern of IEG induction is observed across brain structures (Mukherjee et al., 2018). This transcriptional code was characterized addressing a handful of transcripts within bulk tissue measurements, warranting a comprehensive study of the induced gene expression programs across key structures of the reward circuitry. Here we comprehensively describe gene programs in progressive stages of cocaine experience across multiple brain structures, analyze the spatial and cell-type-specific patterns of IEG expression within prominently recruited brain regions and functionally link induced gene expression to the development of cocaine preference.

Taking an unbiased approach to the identification of the cellular and molecular modifications underlying the development of cocaine-elicited behaviors, we analyzed cocaine-induced transcription dynamics across five structures of the reward circuitry. Of these, the most prominent induced gene programs were identified within the DS. Addressing the spatial segregation of these transcriptional programs within the DS (studying 759,551 individual cells by multiplexed single molecule fluorescence in-situ hybridization), we studied the dynamics of cell-specific recruitment within the two regions engaged by cocaine, the MS and VLS. While both D1R^+^ and D2R^+^ neurons in the MS were engaged transcriptionally throughout the development of cocaine sensitization, we observed dynamic transcriptional recruitment of a cluster of D1R^+^ neurons in the VLS. The IEG *Egr2*, which is the gene we find to be most robustly induced following cocaine experience, serves as a prominent marker for these VLS ensembles. We therefore addressed the function of VLS Egr2^+^ ensembles in the development of cocaine reward, as well the role of Egr2-transcriptional complexes in VLS neurons in the context of cocaine reward. Our results identify the VLS as a hub of dynamic transcriptional recruitment by cocaine, and define a role for Egr2-dependent transcriptional regulation in VLS D1R^+^ neurons in the development of cocaine reward.

## Results

### Characterization of transcriptional dynamics in the reward circuitry during the development of behavioral sensitization to cocaine

In order to characterize brain-wide gene expression programs corresponding to the development of psychostimulant sensitization, we exposed mice to cocaine (20 mg/kg, IP), acutely, or repeatedly (5 daily exposures), as well as to challenge cocaine (acute exposure following 21 days of abstinence following repeated cocaine). We then profiled transcription (applying 3’-RNAseq) within key brain structures of the reward circuitry (limbic cortex=LCtx, nucleus accumbens=NAc, dorsal striatum=DS, amygdala=Amy, lateral hypothalamus=LH; see Figure 1–figure supplement 1 for the definition of individual brain tissue dissected and Figure 1–figure supplement 2 for a description of the samples sequenced) at 0 (not exposed to cocaine on day of sample collection), 1, 2 & 4 hours post cocaine exposure (Figure 1A, B). Mice exhibited a characteristic increased locomotion upon acute exposure to cocaine, further increasing following repeated exposure and maintained after abstinence and challenge re-exposure, typical of cocaine-induced locomotor sensitization (Figure 1B, ANOVA p<0.0001).

**Figure 1:**
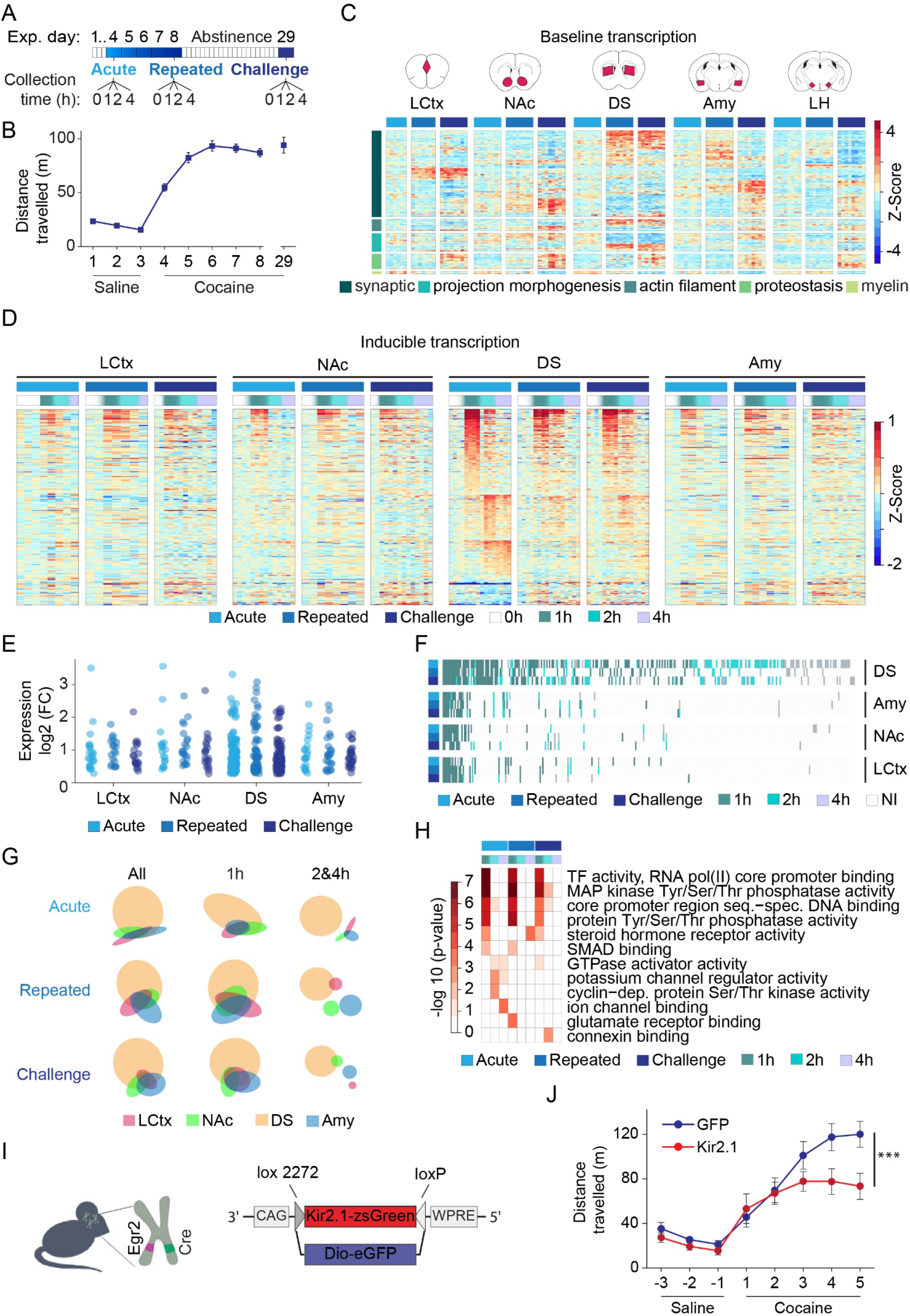
Transcriptional profiling resolves the dynamics of cocaine-induced gene expression within major nodes of the reward circuitry. (**A**) Scheme describing the cocaine sensitization paradigm, and time points (0,1,2,4 hrs) at which samples were obtained for analysis of gene expression following acute (0 = cocaine naïve); repeated (5^th^ exposure to cocaine; 0 = 24 hrs following 4^th^ exposure); and challenge exposures (acute exposure following 21 days of abstinence from repeated exposure; 0 = abstinent mice). **(B)** Locomotor sensitization to cocaine (20 mg/kg i.p.; Days 1-3 n=58; days 4 n=51; day 8 n=30; day 29 n=15). **(C)** Baseline shifts in expression of genes associated with categories of neuroplasticity following repeated cocaine exposure and abstinence. Heatmap depicting fold-change (FC) of differentially-expressed genes (normalized to cocaine naive samples and Z-scored per gene), with rows corresponding to individual genes, clustered according to annotation of biological function on Gene Ontology (p<0.05 FDR corrected). Columns correspond to individual mice of acute (=azure); repeated (=blue); challenge (=navy) cocaine; n = 6 - 8 samples in each group across brain structures (LCtx = limbic cortex, NAc = nucleus accumbens, DS = dorsal striatum, Amy = amygdala and LH = lateral hypothalamus). Genes were selected from a subset of samples which were sequenced together (Fig. 1 – figure supplement 3A) and plotted here across all available samples. **(D)** Heatmaps depicting expression of inducible genes. Data was normalized to 0h of relevant cocaine experience, log-transformed and clustered by peak expression (selected by FC>1.2 and FDR corrected p<0.05, linear model followed by LRT, see methods). Columns correspond to individual mice (0,1,2,4 hrs following acute, repeated vs challenge cocaine; see adjacent key for color coding) across LCtx, NAc, DS and Amy. n= 2-4 samples for individual time points of a cocaine experience within a brain nucleus. **(E)** Dot plots represent the peak induction magnitude of genes induced in the LCtx, NAc, DS and Amy following acute, repeated and challenge cocaine. **(F)** Heatmap addressing the conservation of gene identity and peak induction time. Induced genes are color coded by their time point of peak induction. **(G)** Venn diagrams represent overlap of the genes induced in each brain nuclei following different cocaine experiences (all: 1 & 2 & 4 hrs; early: 1h; late: 2h and 4h time points). **(H)** DEGs induced within the DS are enriched for GO-terms associated with signaling and transcription at 1hr, diversifying to regulators of cellular function and plasticity at later times. Heatmap represents significantly enriched GO-terms (*p* < 0.05, Bonferoni corrected), graded according to *p*-value. **(I)** Illustration of the conditional approach implemented to achieve constitutive inhibition of cocaine responsive neuronal assemblies in the dorsal striatum (DS *Egr2*^+^ neurons), applying double-floxed inverse open reading frame (DIO) viral constructs to express Kir2.1 (vs eGFP controls) in *Egr2*^+^ striatal cells. **(J)** Constitutive inhibition of DS Egr2^+^ neurons attenuates cocaine sensitization. n= 3 mice in each group (***p < 0.0001, Mixed effects linear model; see Supplementary file 6).

### Repeated cocaine administration and abstinence induce prominent transcriptional shifts across multiple brain regions

Experience impacts gene transcription at multiple timescales (Clayton et al., 2020; Mukherjee et al., 2018; Nestler and Lüscher, 2019; Rittschof and Hughes, 2018; Sinha et al., 2020; Yap and Greenberg, 2018). Whereas the expression of inducible genes peak and decay on a time scale of minutes-to-hours following stimulation, baseline shifts in brain-wide gene expression programs are also observed following more prolonged periods (days-weeks) (Clayton et al., 2020) presumably maintaining the modified behavioral output (Sinha et al., 2020).

We initially focused on baseline shifts in gene expression, comparing naïve mice (never exposed to cocaine) to mice exposed repeatedly to cocaine, as well as to mice following 21 days of abstinence after repeated cocaine exposure (Figure 1C; Figure 1–figure supplement 3A; refer to Supplementary file 2 for list of differentially expressed genes and normalized counts). Differentially expressed genes (DEGs) included both upregulated and downregulated genes across all brain regions analyzed, with prolonged abstinence driving the most robust shifts in expression (Figure 1–figure supplement 3B, C). While repeated cocaine induced gene expression shifts were prominent in the DS, abstinence-induced changes were more prominent in the NAc and LCtx (Figure 1–figure supplement 3C). KEGG-analysis demonstrated that DEGs were enriched for synaptic genes and disease pathways (Figure 1–figure supplement 3D). To provide insight into the cellular mechanisms affected by repeated drug exposure and abstinence, we implemented Gene Ontology (GO term) enrichment analysis. (Figure 1C, Figure1–figure supplement 4, see Supplementary file 3 for definition of clusters and DEGs included within them). Gene clusters associated with synaptic plasticity, myelin, and proteostasis demonstrated shifts in expression across multiple brain structures, whereas a cluster of genes associated with structural plasticity appeared more specific to striatal structures (DS and NAc). Noteworthy gene clusters that displayed modified expression were involved in cell-cell communication, glutamate induced plasticity, synaptic vesicle formation, transport, and fusion, actin filament components, projection morphogenesis. Notably, protein folding were coordinately upregulated across structures, while myelin components were coordinately downregulated (Figure 1-figure supplement 4). These results exemplify the dramatic shifts of transcription occurring in the brain in response to repeated cocaine exposure, potentially supporting the maladaptive neuroplasticity driving drug addiction (Bannon et al., 2014; Lull et al., 2008).

### Transcriptional profiling illustrates dynamic recruitment of the striatum during the development of behavioral sensitization to cocaine

Inducible transcription supports the development of plastic changes following psychostimulant experience (Han et al., 2019; Nestler and Lüscher, 2019). We therefore assessed the inducible transcription response 1, 2, or 4 hours following acute, repeated or challenge cocaine exposure, observing robust IEG induction across all brain structures studied (Figure 1D). The largest number of induced genes, as well as the most prominent fold induction levels were found in the DS (Figure 1D, E; refer to Supplementary file 4 for the identities of genes induced in each structure and cocaine condition).

To what extent do the transcription programs induced in the different structures share common attributes? To query the overlap in the identity of genes induced and their temporal induction patterns following the different regiments of cocaine exposure, we graphed the induced genes, color-coding them according to their time of peak induction (1, 2 or 4 hrs following cocaine) (Figure 1F). Thus, for example, if a gene was commonly induced across structures with a peak at 1h across cocaine regiments, this would be evident as a contiguous vertical green line. This graph reveals some aspects of the logic of these inducible transcription programs, whereby: A) genes induced following the different cocaine regiments largely maintain the same temporal structure. Therefore, if the peak induction of a given gene was observed at a defined time-point in one program, its peak induction time was maintained across other programs; B) following repeated cocaine exposure, we observe a substantial dampening of the transcriptional response in the DS, which recovers following cocaine challenge, recapitulating a significant proportion of the acute cocaine gene program; C) all gene programs largely represent subcomponents of the program induced by acute cocaine in the DS. We further visualized the overlap in the identity of genes induced in the different structures using venn diagrams (Figure 1G), illustrating that the overlap stems principally from the immediate component of the transcriptional program (peaking at 1hr following cocaine), while transcripts induced at 2 or 4 hrs following cocaine diverged between structures. Focusing on the most robust programs, induced in the DS, we found that gene clusters enriched at the 1 hr time points are related primarily to transcriptional regulation and synapse-to-nucleus signal transduction, while clusters related to modification of neural morphology and function were enriched at later time points (Figure 1H; refer to Supplementary file 5). Taken together, these results highlight robust transcriptional adaptations in the DS, suggesting that it is a major hub of cocaine-induced plasticity. Furthermore, our results illustrate the utilization of a conserved set of genes during the early wave of transcription following experience, followed by divergence of subsequent transcription, possibly to support region-specific mechanisms of plasticity (Hrvatin et al., 2018; Walker et al., 2018).

### Striatal *Egr2*-expressing ensembles contribute to cocaine sensitization

Recently, we have shown that salient experiences are represented in the mouse brain by unique patterns of gene expression. Thus, the induction pattern of a handful of genes was found to be sufficient to decode the recent experience of individual mice with almost absolute certainty. Of these, the IEG whose expression contributes most towards classification of the recent experience of individual mice is *Egr2* (Mukherjee et al., 2018)*. Egr2* is, furthermore, the gene most robustly induced by cocaine in the dorsal striatum [(Gonzales et al., 2020; Mukherjee et al., 2018); Supplementary file 4], and is a sensitive indicator of cocaine-engaged striatal cell assemblies (Gonzales et al., 2020). Transgenic approaches based on IEG expression support genetic access to neuronal assemblies recruited by experience (Cruz et al., 2013; DeNardo and Luo, 2017; Hope, 2019; Minatohara et al., 2016; Sakaguchi and Hayashi, 2012; Tonegawa et al., 2015). To address the potential role of dorsal striatal assemblies expressing *Egr2* in the development of behavioral sensitization to cocaine, we utilized transgenic Egr2-Cre knock-in mice, expressing Cre recombinase under the regulation of the Egr2 promoter (Voiculescu et al., 2000). In these mice, we inhibited the activity of Egr2-expressing neuronal ensembles, by AAV-mediated expression of a CRE-dependent inward-rectifying K^+^-channel (DIO-Kir2.1) (Atlan et al., 2018; Rothwell et al., 2014; Terem et al., 2020), aiming to transduce a large proportion of the DS. Control mice were transduced with DIO-GFP (DS-Egr2^GFP^) (Figure 1I; infections exhibited a bias for the medial aspect of the striatum; for documentation of sites of infection see Figure 1–figure supplement 5). While control mice (DS-Egr2^GFP^) developed stereotypical behavioral sensitization to cocaine (Figure 1J), experimental mice, in which striatal *Egr2*^+^ neurons had been constitutively inhibited, developed attenuated behavioral sensitization (Figure 1J; p<0.0001, ANOVA). The distinction between the two groups of mice became apparent on Day 3 of cocaine administration, segregating further on Day 4 and 5 (see Supplementary file 6 for detailed statistics). This result demonstrates that the activity of Egr2-expressing neuronal assemblies in the DS is functionally relevant for the development of cocaine-induced locomotor sensitization.

### IEG induction in sub-compartments of the DS is influenced by the history of cocaine exposure

Our observation of dynamics within the transcriptional response to repeated cocaine exposure in the DS (Figure 1) motivated us to address the cellular basis and spatial distribution of this transcriptional plasticity. Recently, we reported, using single molecule fluorescence in-situ hybridization (smFISH), region-specific rules governing the recruitment of striatal assemblies following a single acute exposure to cocaine (Gonzales et al., 2020). We now revisited this spatial analysis, applying smFISH to study the striatal distribution of the IEGs *Arc*, *Egr2*, *Fos* and *Nr4a1* throughout the development of cocaine sensitization [Figure 2; Figure 2–figure supplement 1, 2; (Gonzales et al., 2020)]

**Figure 2:**
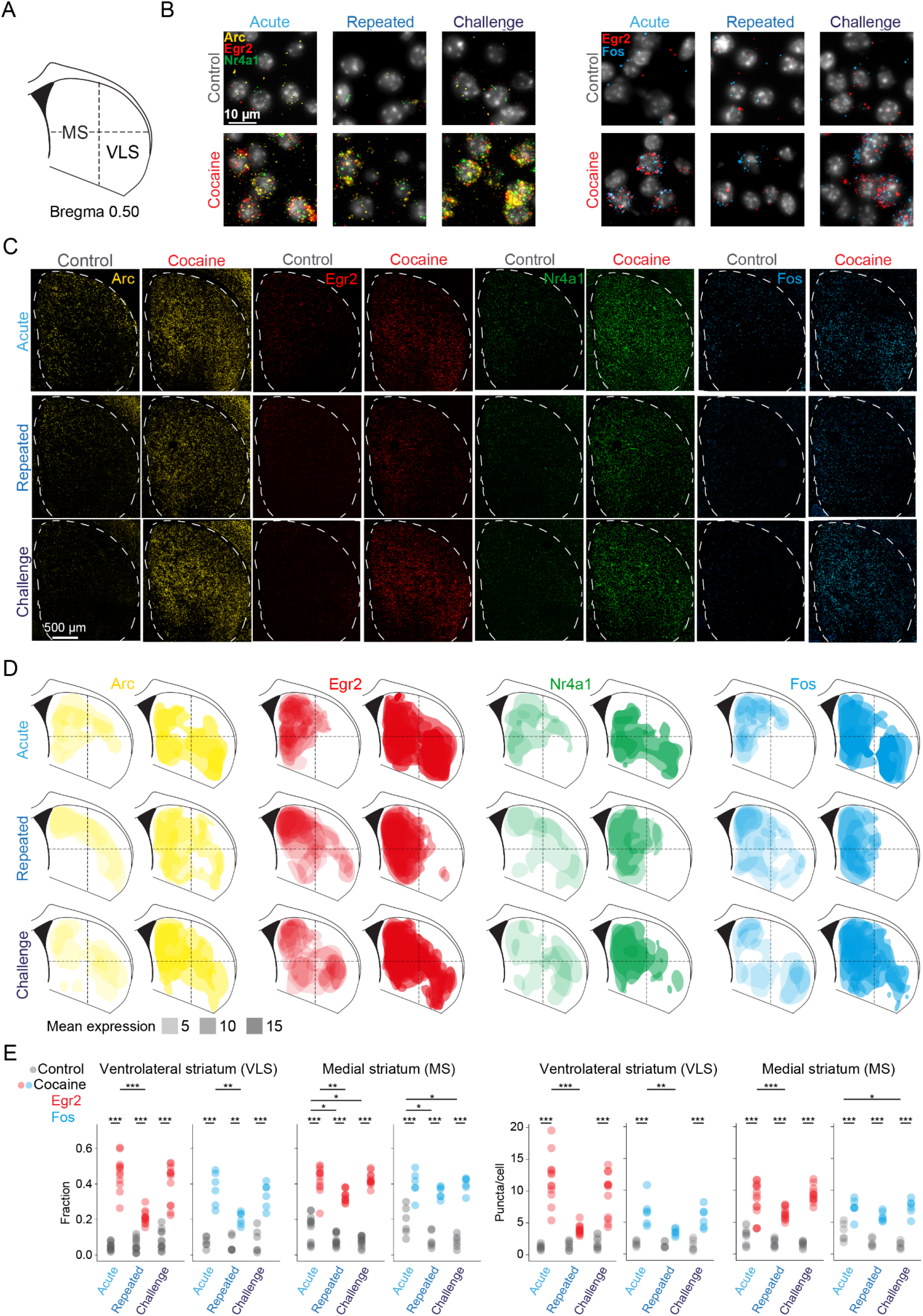
Dynamic IEG induction in subregions of the striatum accompany the development of cocaine sensitization. (**A**) Representative coronal section of the dorsal striatum (DS) (∼0.52 ± 0.1 mm from Bregma) assayed by multicolor smFISH for cocaine-induced IEG expression. (**B**) Representative images of multicolor smFISH analysis of *Arc*, *Egr2*, *Nr4a1* and *Fos* expression following acute, repeated and challenge cocaine exposures (40X magnification). (**C**) Spatial IEG expression patterns in the DS. Representative images of multicolor smFISH analysis of *Arc*, *Egr2*, *Nr4a1* and *Fos* expression. **(D)** Cocaine experiences induce distinct spatial patterns of IEG expression. Two-dimensional kernel density estimation was used to demarcate the regions with maximal density of high expressing cells for each IEG. Color code for probes: *Arc*-yellow, *Egr2*-red, *Nr4a1*-green, *Fos*-blue. The opacity of the demarcated areas corresponds to the mean per-cell expression. **(E)** Dot-plots depicting the proportion of suprathreshold cells (fraction) and cellular expression (puncta/cell) of *Egr2*^+^ and *Fos*^+^ in the ventrolateral (VLS) and medial (MS) striatum following acute, repeated and challenge cocaine. \**p*<0.05, \*\**p*<0.005, \*\*\**p*<0.0001, Two-way ANOVA with post hoc Tukey’s test. Refer to Supplementary file 7 for cell numbers.

Addressing an overview of induced expression of these IEGs, we observed robust induction of *Arc*, *Egr2*, *Nr4a1* and *Fos* following acute cocaine exposure, which was dampened following repeated exposure to cocaine and reinstated following a challenge dose of cocaine, in-line with the results described in Figure 1 (Figure 2A-C). To visualize the domains defined by IEG expressing cells, we applied 2D kernel density estimation on striatal sections following repeated and challenge cocaine, and compared these patterns to the expression domains following acute exposure to cocaine. This analysis revealed the dynamic recruitment of two distinct “hotspots” of IEG expression following acute cocaine exposure, corresponding to the medial (MS) and ventrolateral (VLS) aspects of the striatum (Gonzales et al., 2020).

The domain of IEG induction within the MS appeared to be recruited to a largely similar extent following acute, repeated and challenge cocaine. In stark contrast, the recruitment of the prominent IEG expression domain in the VLS following acute cocaine exposure was dampened drastically following repeated cocaine exposure, re-emerging following cocaine challenge (Figure 2D). These results are quantified in Figure 2E, demonstrating the dampened VLS response (Mean difference (cocaine-control): **VLS *Egr2***: Acute: 41±8% expressing, 11±3 puncta/cell; Repeated: 15±8%, 3±3 puncta/cell; Challenge: 31±8%, 8±3 puncta/cell; **VLS *Fos***: Acute: 30±10%, 5±2 puncta/cell; Repeated: 14±10%, 2±2 puncta/cell; Challenge: 25±10%, 4±2 puncta/cell; All number are mean ± CI, for statistics refer to Supplementary file 6). In the MS, in contrast, a substantially smaller effect of repeated cocaine exposure was observed (**MS *Egr2***: Acute: 27±6%, 6±2 puncta/cell; Repeated 24±6%, 5±2 puncta/cell; Challenge: 34±6%, 8±2 puncta/cell; **MS *Fos***: Acute: 20±9%, 4±2 puncta/cell; Repeated 26±9%, 4±2 puncta/cell; Challenge: 32±9%, 6±2 puncta/cell) (Figure 2E). A similar trend was evident for the expression of *Arc* and *Nr4a1* in the VLS vs. the MS (Figure 2–figure supplement 1A, B). Notably, the expression of IEGs was highly correlated within individual cells, demonstrating that the expression of the different IEGs define overlapping populations of neurons responsive to the cocaine experiences studied. Once recruited by cocaine, neurons commit to co-expression of multiple IEGs to virtually identical levels (Figure 2–figure supplement 2; see Supplementary file 6). These data demonstrate the coherent co-expression of multiple IEGs within striatal assemblies during the development of behavioral sensitization to cocaine, likely in order to support mechanisms of long-term plasticity within these ensembles. In sum, the history of cocaine experience is reflected in a differential manner in the transcriptional recruitment of striatal sub-compartments, dampening drastically in the VLS following repeated exposure, while exhibiting continuous maintenance in the MS.

### The IEG response in VLS *Drd1*^+^-SPNs is selectively dampened following repeated cocaine

Striatal Drd1^+^-neurons are implicated in promoting actions, while *Drd2*^+^-neurons are implicated in the refinement and tempering of action selection (Bariselli et al., 2019). Differential IEG induction in *Drd1* vs *Drd2* expressing SPN ensembles is expected to shed light on the relative contribution of plasticity within each cell type to the development of cocaine behaviors. We have previously reported that acute exposure to cocaine induces *Egr2* expression in both *Drd1*^+^ and *Drd2*^+^ neurons in the MS, while selectively inducing *Egr2* expression in *Drd1*^+^-neurons in the VLS (Gonzales et al., 2020). Extending this analysis to repeated and challenge cocaine exposures (Figure 3A, B, Figure 3 – figure supplement 1) and with additional IEGs, we observed robust dampening of the induction of *Egr2* and *Fos* in VLS *Drd1*^+^ neurons following repeated exposure to cocaine, which regains the prominent *Drd1*-specificity of induction following cocaine challenge. In contrast, while subtle dampening was also observed in the MS, *Egr2* and *Fos* expression maintained consistent correlation to *Drd1*- and *Drd2*- expression throughout acute, repeated and challenge cocaine exposures (Figure 3A, B, Figure 3 – figure supplement 1A-C; for reference of *Drd1* and *Drd2* levels in MS and VLS see Figure 3 – figure supplement 2, Supplementary file 6 for stats). Thus, a recent history of repeated cocaine exposure results in a dampening of transcriptional recruitment in the DS, which can largely be attributed to the selective dampening of induction within VLS *Drd1*^+^-neurons. This specialization in transcriptional plasticity likely underlies differential roles of the striatal subregions and cells within them in supporting behavioral modification induced by cocaine experience.

**Figure 3:**
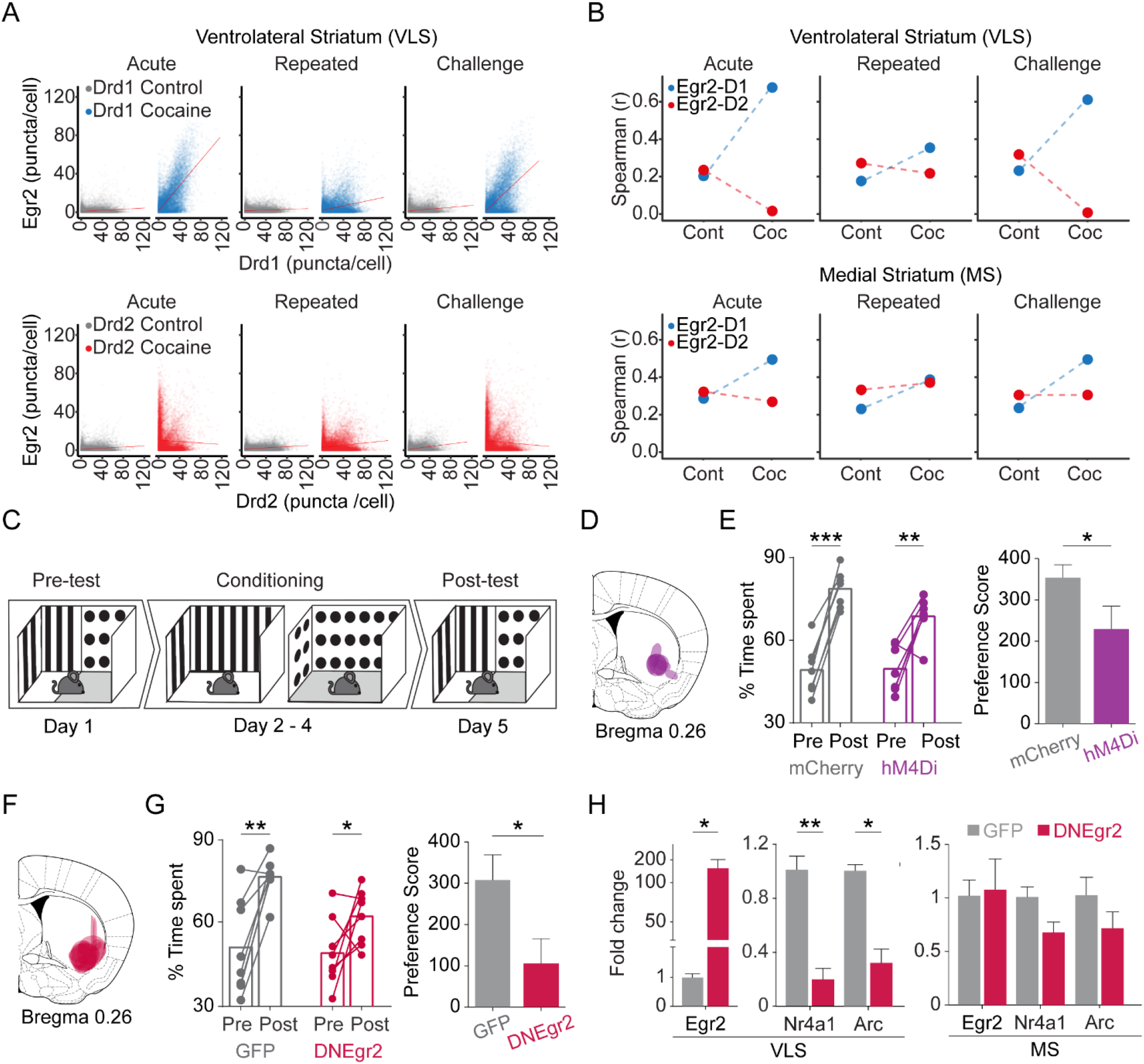
Induction of *Egr2* in VLS Drd1+ neurons contributes to the acquisition of cocaine reward. **(A, B)** The *Drd1*^+^-specific IEG response in the VLS is dampened following repeated exposure to cocaine. **(A)** Scatter plots show cellular *Egr2* expression with *Drd1* or *Drd2* expression (puncta/cell) within individual cells. n=6 sections from 3 mice for each condition (grey-0h for either *Drd1* or *Drd2* combination, and blue or red – for *Drd1* or *Drd2* combination, respectively, 1h following cocaine experience). Refer to Supplementary file 6 for detailed statistics. (**B**) Spearman correlation plots showing acute induction of *Egr2* is correlated with *Drd1* expression in the VLS, dampened following repeated exposure and re-emerges following challenge exposure. In the MS, *Egr2* expression is consistently correlated to both *Drd1* and *Drd2* expression for acute, repeated and challenge exposures. (**C**) Scheme of experimental paradigm for testing conditioned-place preference (CPP) for cocaine. (**D**) Summary of expression domains of AAV-DIO-h4MDi in Egr2-CRE mice. **(E)** Chemogenetic inhibition of VLS-Egr2 expressing neurons during conditioning attenuates the development of cocaine CPP. Egr2-CRE animals were stereotactically transduced with AAV-DIO-mCherry (VLS-Egr2^mCherry^) or AAV-DIO-hM4Di-mCherry (VLS-Egr2^hM4Di^) and following recovery subjected to cocaine CPP conditioning 30 minutes following exposure to CNO. Left panel represents change in % time spent on the cocaine paired side before and after conditioning for individual animals and the mean (paired t-test), while right panel (bar graphs) displays the mean preference score (time spent on the drug paired side of the final – first test day; unpaired t-test). Both groups developed CPP (paired t-test), while VLS-Egr2^hM4Di^ mice displayed a lower preference score compared to VLS-Egr2^mCherry^ controls (unpaired t-test). n = 7 mice in each group. \**p*<0.05, \*\**p*<0.01, \*\*\**p*<0.005. (**F**) Summary of expression domains of AAV-DN-Egr2. **(G)** Disruption of *Egr2* function in the VLS inhibits the development of cocaine place preference. Left panel represents change in % time spent on the cocaine paired side before and after conditioning for individual animals and the mean, while bar graphs (right panel) display the mean preference score. Both groups developed CPP (paired t-test), while AAV-DNEgr2 injected displayed attenuated CPP in comparison to AAV-GFP injected mice (unpaired t-test). n = 8 mice in each group. \**p*<0.05, \*\**p*<0.01, \*\*\**p*<0.005**. (H)** AAV-DN-Egr2 inhibits *Arc* and *Nr4a1* expression in the VLS but not in the MS. Comparison of gene expression in mice injected with AAV-GFP (grey) or AAV-DN-Egr2 (maroon) 24h after the final CPP test. Data represented as mean ± sem. \**p*<0.05, \*\**p*<0.005 unpaired t-test, n= 3 mice in each group.

### VLS *Egr2* transcriptional activity is necessary for the development of cocaine-seeking behavior

The selectivity of Egr2 induction to VLS *Drd1*^+^ neurons suggested a causal role for this neuronal population in supporting cocaine-conditioned behaviors. To address the role of VLS *Egr2*^+^ neurons in cocaine seeking, we performed viral transduction of the Cre-dependent hM4Di DREADD (VLS-Egr2^hM4Di^) or mCherry (control: VLS-Egr2^mCherry^), targeting the VLS of Egr2-Cre knock-in mice (for expression domains see Figure 3D, Figure 3 - figure supplement 3A,B). The selective expression of hM4Di enables inhibition of *Egr2*^+^ neurons upon administration of CNO (10mg/kg; i.p.) (Atlan et al., 2018; Terem et al., 2020). We then trained mice on a CPP paradigm in which we injected CNO (10 mg/kg) 30 min prior to cocaine training sessions (Figure 3C). We found that while both groups developed CPP [paired t-test on % time spent on drug paired side; p_VLS-Egr2mCherry_ < 0.00001, p_VLS-Egr2hM4Di_ < 0.003], mice in which VLS *Egr2*^+^ neurons were inhibited (VLS-Egr2^hM4Di^) displayed attenuated CPP compared to control mice (VLS-Egr2^mCherry^) (Figure 3E; p<0.05). Notably, no differences in locomotion were observed between groups on conditioning or test days (p = 0.93, ANOVA; Figure 3 – figure supplement 3C). We therefore concluded that VLS *Egr2*^+^-expressing neurons contribute to the development of cocaine-seeking behavior, with no obvious impact on locomotor aspects of cocaine-driven behaviors.

Salient experiences in general, and specifically exposure to cocaine, are thought to modify future behavior through induced gene expression responses, leading to stable changes in cell and circuit function (Nestler and Lüscher, 2019; Robison and Nestler, 2011). To assess a potential link between the expression of *Egr2* and cellular plasticity responsible for cocaine-induced modification of behavior, we addressed a causal role for the induction of the *Egr2* gene by cocaine within VLS neurons. To this end, we trained mice on a conditioned place preference paradigm, following viral transduction of GFP (VLS^GFP^) in the VLS, or disruption of Egr2 function by transduction of a dominant-negative (S382R, D383Y) isoform of *Egr2* (VLS^DNEgr2^) in VLS neurons (Figure 3F, Figure 3-figure supplement 4A, B). The dominant-negative mutation of Egr2 disrupts the DNA-binding activity of Egr2, while not interfering with the capacity of the protein to form heteromeric complexes with its natural binding partners, effectively inhibiting transcriptional activation of downstream genes regulated by Egr2 (LeBlanc et al., 2007; Nagarajan et al., 2001). Comparing the development of cocaine conditioned-place preference, we found that both groups of mice developed CPP (Figure 3G, paired t-test on % time spent on drug paired side; p_VLS-GFP_ < 0.001, p_VLS-DNEgr2_ < 0.05). However, VLS^DNEgr2^ mice displayed attenuated CPP in comparison to the VLS^GFP^ mice (Figure 3G, p< 0.05). No differences in locomotion were observed between the groups of mice (p = 0.7, ANOVA; Figure 3–figure supplement 4C). These results assign a functional role to *Egr2* induction, putatively within VLS *Drd1*^+^ neurons, in the development of conditioned place preference to cocaine. To test the effect of the disruption of Egr2 complexes on transcription, we analyzed the expression of *Arc*, *Egr2* and *Nr4a1* in the VLS, MS, NAc and LCtx. In the VLS, we observed the anticipated overexpression of *Egr2* (Figure 3H, p_(Egr2)_ < 0.05, reflecting exogenous expression of the mutant gene), as well as blunted *Arc* and *Nr4a1* expression (Figure 3H, t-test; p_(Arc)_ < 0.01, p_(Nr4a1)_ < 0.005). We did not observe any clear changes between groups in gene expression within other structures, demonstrating the predominantly localized effect of our viral manipulation (Figure 3–figure supplement 4D,E). These results demonstrate a role for cocaine-induced expression of *Egr2* in the VLS in supporting the development of cocaine-seeking, and suggest that inducible transcriptional complexes involving Egr2 are functional in facilitating drug-induced maladaptive plasticity.

## Discussion

Drugs of abuse such as cocaine are known to act on key brain circuits, modifying and biasing the future behavior of an individual toward increased drug seeking. In this study we develop a comprehensive compendium of the transcriptional dynamics induced within key brain regions during the development of cocaine sensitization. We highlight the striatum as a major hub of plasticity, within which we identify differential transcriptional recruitment of neuronal ensembles by cocaine, depending on striatal sub-region, identity of projection neuron and the history of cocaine exposure. Finally, we focus on a prominent cocaine-sensitive IEG, *Egr2*, and show that *Egr2*-expressing (putatively *Drd1*^+^) SPNs in the VLS, and the expression of *Egr2* within them, support drug-seeking behavior and context association to cocaine reward.

Repeated exposure to cocaine, as well as abstinence from drug exposure, produce long lasting functional changes in the reward circuit to drive the maladaptive modification of reinforced behavior (Dong and Nestler, 2014; Everitt, 2014; Gremel and Lovinger, 2017; Hyman et al., 2006; Kelley, 2004; Lüscher, 2016; Lüscher and Malenka, 2011; Nestler, 2013; Russo and Nestler, 2013; Salery et al., 2020; Volkow and Morales, 2015; Wolf, 2016; Zahm et al., 2010). Cocaine-induced alterations in gene expression support the imprinting of such potentially lifelong alterations driven by drug experience (McClung and Nestler, 2008; Nestler, 2002; Nestler and Lüscher, 2019; Salery et al., 2020; Steiner, 2016; Steiner and Van Waes, 2013). In this study, using an unbiased approach to screen gene expression, we resolve the transcriptional landscapes of multiple reward-related brain circuits that characterize distinct cocaine experiences with broad temporal resolution. Our approach allowed us to describe the transcripts modulated at both baselines (i.e. shifts driven following repeated exposure to cocaine or following abstinence from repeated exposure), and following exposure to distinct cocaine experiences (IEG expression).

Baseline transcriptional changes in cortical and basal ganglia structures following defined cocaine regiments have been described previously in both rodents and humans (Bannon et al., 2005, 2014; Eipper-Mains et al., 2013; Freeman et al., 2010; Gao et al., 2017b; Hurd and Herkenham, 1993; Lull et al., 2008; Ribeiro et al., 2017; Walker et al., 2018). Consistent with previous findings, we observed dynamic shifts in baseline gene expression in multiple categories potentially associated with neuronal plasticity (synaptic genes; genes associated projection morphogenesis, actin filament regulation; proteostasis and myelin). Interestingly, genes associated neuronal morphology and synaptic function demonstrated unique patterns of shifts within different brain structures. For example, expression of the genes such as *Vamp*, *Pkrcg*, *Ncdn*, *Camk2b, Shank3, and Syp* were downregulated in the NAc following repeated cocaine exposure, while being upregulated in the DS. Such region-specific shift in gene expression may support circuit-specific structural and functional modifications to cell assemblies (Clayton et al., 2020; Kyrke-Smith and Williams, 2018). Myelin genes (*Plp1*, *Mobp*, *Mbp*, *Mal*, *Pllp*) were downregulated across all structures studied (LCtx, Amy, NAc, DS, LH), initially decreasing following repeated cocaine exposure, and further decreasing following abstinence, across all experimental mice. Conversely, genes associated with proteostasis (e.g. chaperones such as members of the CCT, Hsp40, Hsp70, and Hsp90 complexes) demonstrated concerted upregulation across structures, occurring following cocaine abstinence. Notably, similar changes in myelin genes and genes associated with proteostasis are consistent in both human and rodent studies, but have not been functionally interrogated (Albertson et al., 2004; Bannon et al., 2014; García-Fuster et al., 2012; Johnson et al., 2012; Kovalevich et al., 2012; Lull et al., 2008; Narayana et al., 2014). Future investigation into the features of cocaine experience-related transcriptome is anticipated to provide targets for intervention, potentially supporting the reversal of brain function to a “cocaine-naive” state.

IEG expression is well accepted to be the substrate for long term modulations supporting memory formation (Alberini, 2009; Alberini and Kandel, 2015). Although cocaine induced IEG expression has been extensively characterized in rodents (Caster and Kuhn, 2009; Gao et al., 2017b; Guez-Barber et al., 2011; Moratalla et al., 1996; Piechota et al., 2010; Robison and Nestler, 2011; Savell et al., 2020; Steiner, 2016; Valjent, 2006; Zahm et al., 2010), these studies were mostly limited in the number of genes analyzed, and restricted to isolated brain structures following specific drug regimes. Addressing the cocaine-induced transcriptome, we observed transcriptional recruitment of the LCtx, Amy, NAc and DS, of which, the DS was most prominent. Furthermore, the immediate-early transcriptional programs induced across other tissues largely comprised of sub-components of the programs induced in the DS. What does this imply? We propose that the overlapping fraction of induced genes are representative of the “core transcriptome” that is consistently induced across all structures or cell types and only vary in the magnitude of their expression (Hrvatin et al., 2018; Tyssowski et al., 2018). This core component predominantly corresponds to signaling molecules and transcriptional regulators (the genes common across most programs are *Arc, Arl4D, Btg2, Ddit4, DUSP1, Egr2, Egr4, Fos, FosB, JunB, Nr4A1, Per1 & Tiparp*), likely responsible for transforming the inducing signal into instructions for implementation of appropriate synaptic, cellular and circuit specific plasticity mechanisms by “effector” genes, induced in a delayed transcriptional wave, corresponding to the significantly diversified gene response at 2,4h following cocaine (Amit et al., 2007; Clayton et al., 2020; Gray and Spiegel, 2019; Hrvatin et al., 2018; Mukherjee et al., 2018; Tyssowski et al., 2018; Yap and Greenberg, 2018).

What might be the role of the transcriptional induction in the DS and its subsequent dampening? It is becoming more broadly accepted that IEG induction serves to support long-term plasticity [rather than simply indicate activity (Chandra and Lobo, 2017; Clayton, 2000; Clayton et al., 2020; Mukherjee et al., 2018; Tyssowski and Gray, 2019)]. Therefore, it is likely that this transcriptional induction serves to support plasticity within these circuits. The MS is defined as the ‘associative striatum’, and is associated with goal-directed behaviors, as well as defining the vigor of locomotor actions (Balleine and O’Doherty, 2010; Balleine and Ostlund, 2007; Kravitz et al., 2010; Lipton et al., 2019; Nonomura et al., 2018). We propose that the cocaine-driven locomotor response may be mediated by the balanced and maintained transcriptional induction within Drd1/Drd2 SPNs in the MS. Furthermore, we observed a dampened development of behavioral sensitization to cocaine upon disrupting the action of cocaine-responsive neuronal ensembles in the DS that were captured through their expression of the IEG *Egr2*, suggesting that recruitment of IEG-expressing striatal SPNs is necessary for the appropriate development of cocaine-induced locomotor sensitization. The lateral ‘sensori-motor’ striatum is strongly associated with habit formation and compulsive drug seeking (Lipton et al., 2019; Yin et al., 2004; Zapata et al., 2010). Moreover, The VLS receives selective sensorimotor afferents mapped to upper limb and orofacial cortical regions. Interestingly, behavioral stereotypies, primarily upper-limb and orofacial, arise upon psychostimulant exposure (Karler et al., 1994; Murray et al., 2015; Schlussman et al., 2003), and orofacial stereotypies have been induced following selective infusion of psychostimulants to the VLS (Baker et al., 1998; Delfs and Kelley, 1990; Rebec et al., 1997; White et al., 1998). It is intriguing to consider the possibility that recruitment of plasticity mechanisms within VLS Drd1^+^ neurons supports the increased propensity to engage in orofacial stereotypies, while the subsequent dampening of cocaine-induced transcription within these neurons may support the ‘canalization’ of this limited action repertoire, at the expense of a broader behavioral repertoire. This topic will form the basis for future investigation.

Infusion of psychostimulants into the VLS has been shown to promote operant reinforcement and conditioned-place preference, implicating it in reward and reinforcement (Baker et al., 1998; Kelley and Delfs, 1991). In order to query the role of the VLS IEG-expressing ensembles in the development of cocaine context association, we inhibited the activity of VLS Egr2^+^ neurons by conditional expression of Kir2.1, and observed an attenuation of the development of CPP. To directly investigate a role for VLS IEG action in the development of CPP, we expressed a dominant-negative isoform of Egr2 (in which the DNA-binding domain was inactivated) in the VLS, and also observed an attenuation of CPP development. We therefore provide the first functional implication of the VLS in cocaine reward, and provide a description of the cellular dynamics of its recruitment during the development of drug reward. The development and execution of drug-seeking behavior is heavily context dependent (Calipari et al., 2016; Crombag, 2002; Crombag et al., 2008; Crombag and Shaham, 2002; Cruz et al., 2014; Lee, 2006; Rubio et al., 2015). Our results suggest that the transcriptional recruitment of VLS Drd1^+^ neurons following acute cocaine exposure is instrumental for cocaine seeking. Potentially, the dampening of sensorimotor VLS IEG induction following repeated cocaine further serves to ‘cement’ the initial context association, limiting behavioral flexibility and the capacity to revert context association, exacerbating the impact of contextual cues on drug seeking behavior (Calipari et al., 2016; Crombag and Shaham, 2002; Gipson et al., 2013; Hyman, 2005; Phillips et al., 2003; Shaham et al., 2003; Volkow et al., 2006).

We further introduce *Egr2* as a molecular player, acting within *Drd1*^+^ neurons of the VLS to promote the development of drug seeking. Previous studies have showed that Egr2 is crucial for normal hindbrain development, peripheral myelination, humoral immune response, and implicated in diseases such as congenital hypomyelinating neuropathy, Charcot–Marie-Tooth disease, Dejerine–Sottas syndrome, as well as Schizophrenia (Boerkoel et al., 2001; De and Turman, Jr., 2005; Li et al., 2019; Morita et al., 2016; Okamura et al., 2015; Svaren and Meijer, 2008; Topilko et al., 1994; Warner, 1999; Warner et al., 1998; Wilkinson, 1995; Yamada et al., 2007). In the central nervous system, *Egr2* has been shown to be induced by seizure activity, kainic acid injection, following LTP-inducing stimuli in hippocampal neurons, as well as following administration of several groups of drugs such as methamphetamine, cocaine, heroin, and alcohol (Gao et al., 2017a; Gass et al., 1994; Imperio et al., 2018; López-López et al., 2017; Mataga et al., 2001; Rakhade et al., 2007; Saint-Preux et al., 2013; Worley et al., 1993). However, its role in encoding memory or drug-induced behavior remained unresolved. Our findings show that the activity of *Egr2* is required for encoding drug-induced memory, and demonstrate that an additional member of the Egr family, alongside *Egr1* and *Egr3,* plays a pivotal role in drug-induced plasticity (Bannon et al., 2014; Chandra et al., 2015; Moratalla et al., 1992; Valjent, 2006). Furthermore, we describe that *Egr2* expression can be reliably utilized to target specific cocaine responsive ensembles in different subregions of the striatum and probe their involvement in driving distinct aspects of cocaine behavior.

In sum, our study provides a comprehensive description of brain-wide transcriptional dynamics, as well as spatial dynamics of SPN-specific IEG recruitment during the development of cocaine sensitization. Furthermore, our results demonstrate the role of striatal IEG-expressing ensembles in the development of cocaine sensitization and contextual association of reward, as well as the role of IEG induction in VLS Drd1^+^ neurons in cocaine reward. Future work will address the mechanisms supporting cell-type specificity of transcriptional induction, as well as the role of IEG-mediated plasticity mechanisms in VLS-dependent stereotypy and context association.

## STAR★Methods

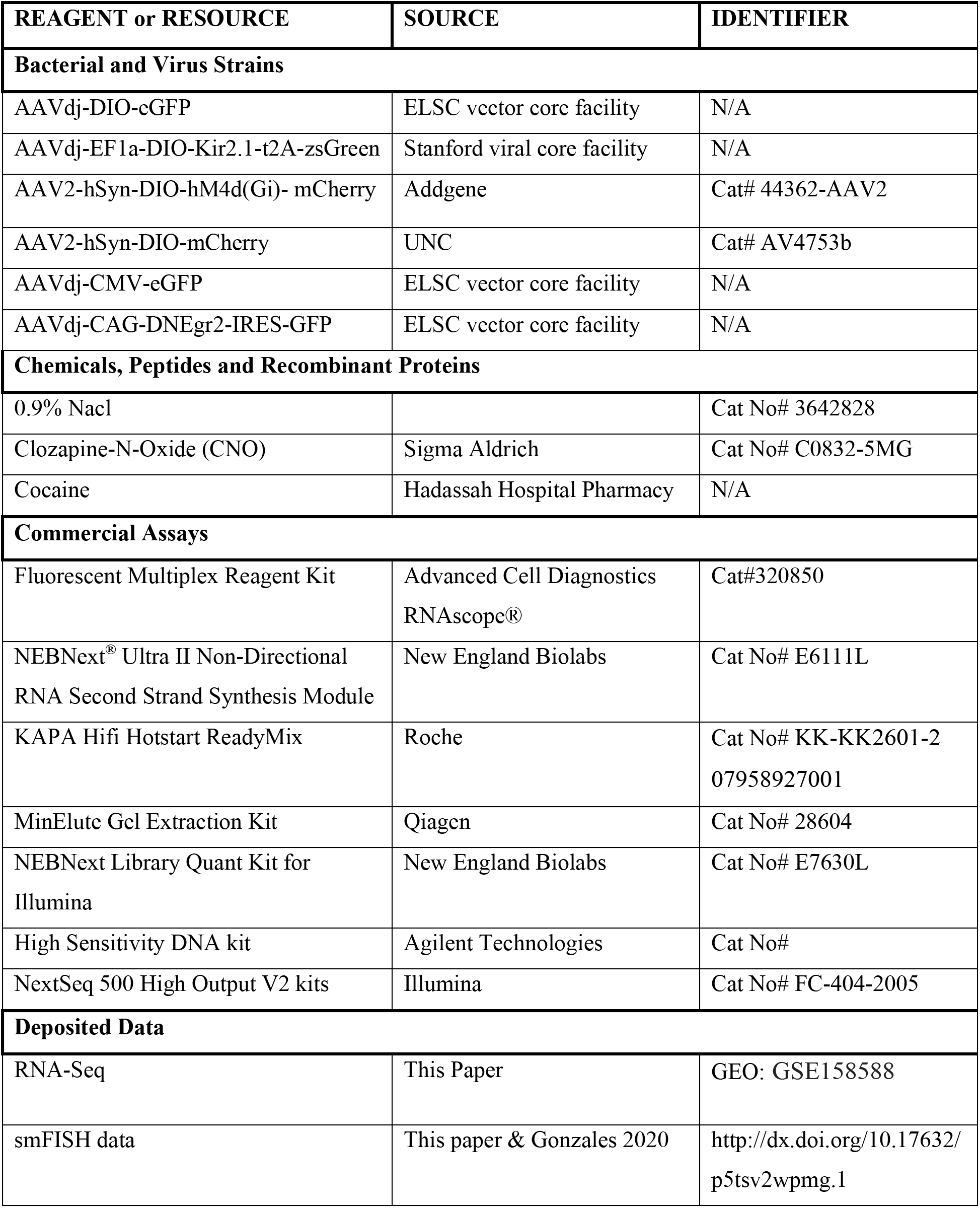
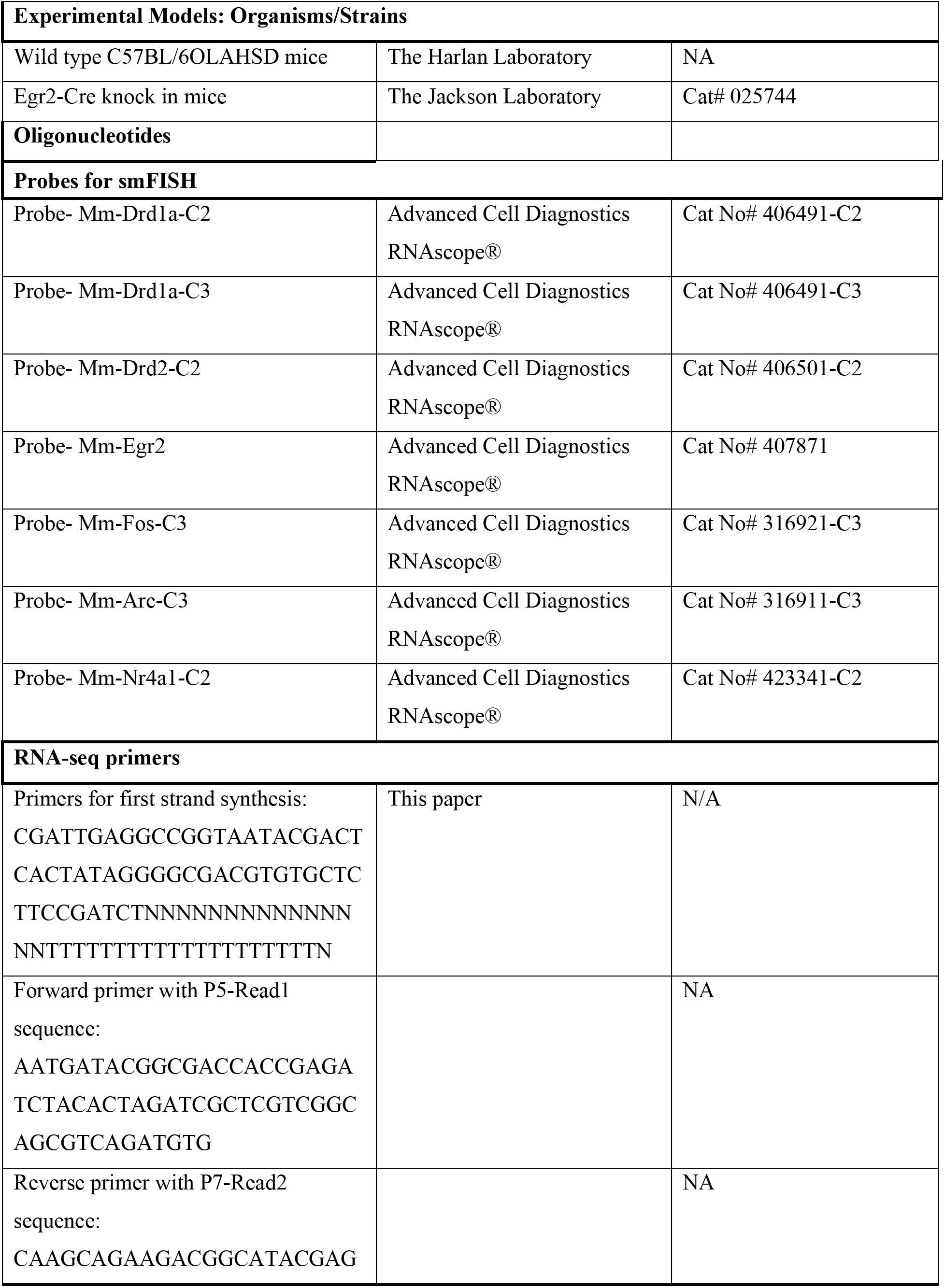
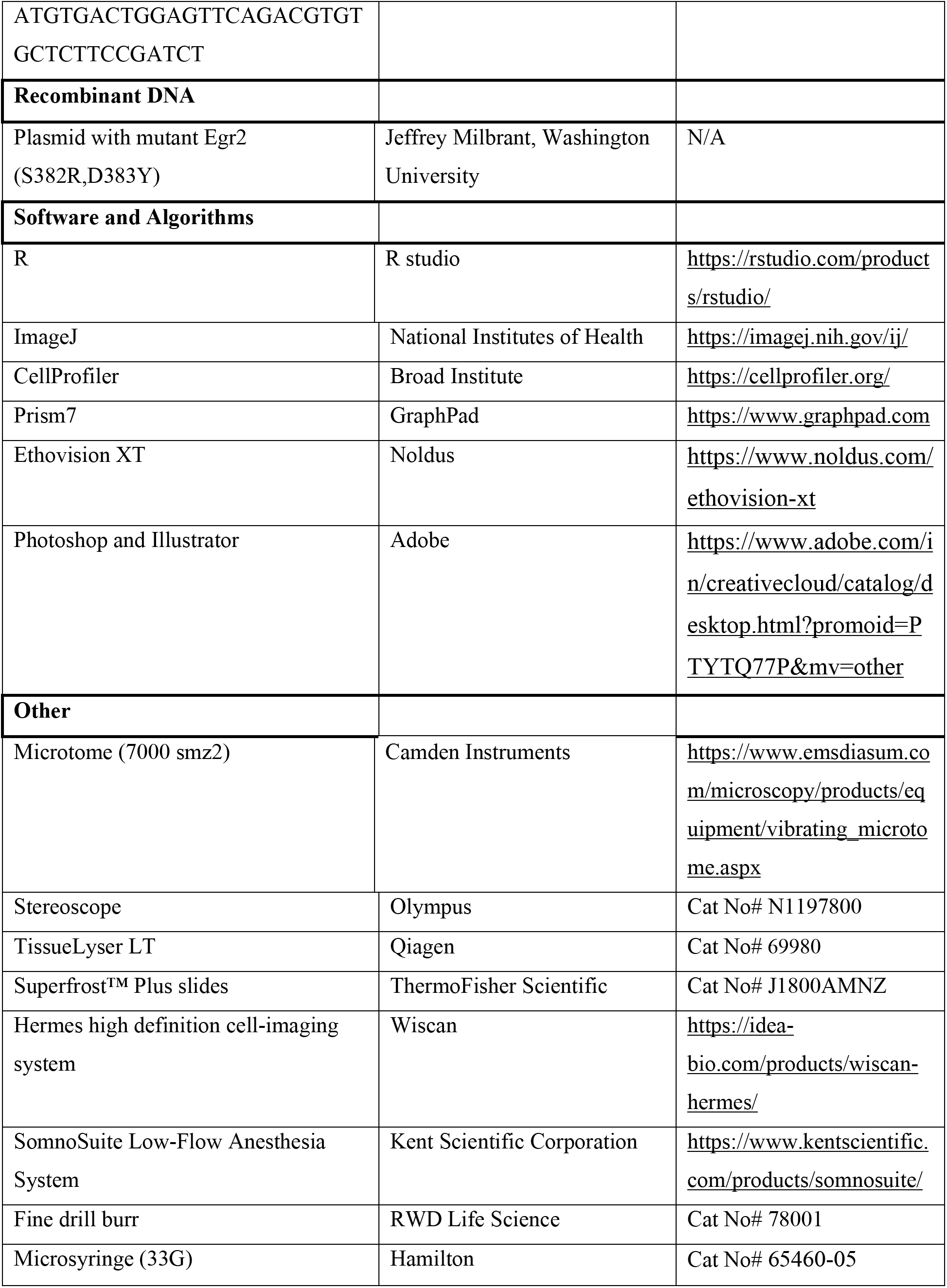
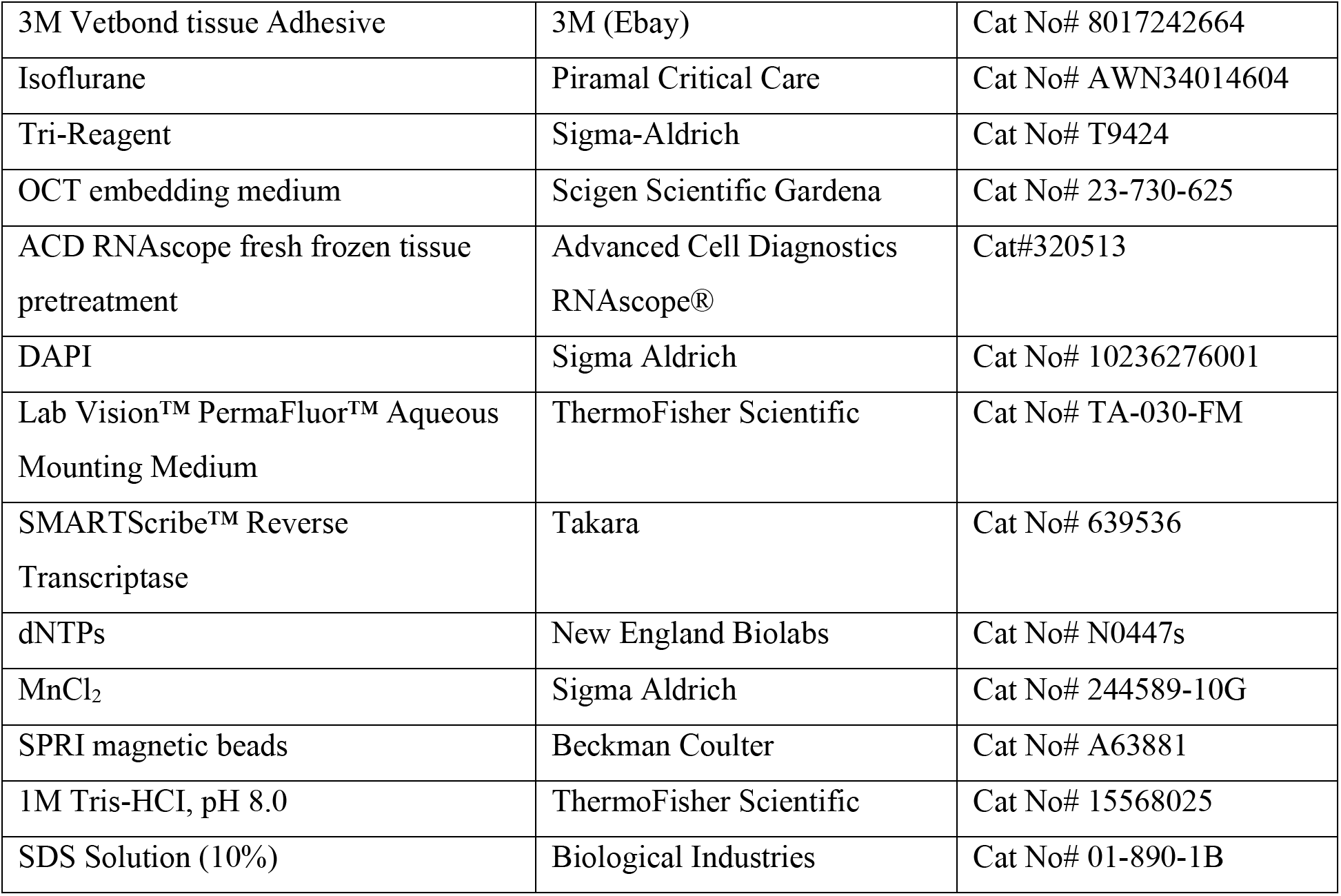
Key resource Table:

### Lead Contact and Materials Availability

Further information and requests for resources should be directed to and will be fulfilled by the Lead Contact, Ami Citri. This study did not generate new unique reagents.

### Experimental models and subject details

Male C57BL/6OLAHSD mice used for RNA-sequencing and single molecule-FISH analysis following cocaine sensitization were obtained from Harlan Laboratories, Jerusalem, Israel. Transgenic Egr2-Cre knock-in mice were obtained from Jackson Laboratories. All animals were bred at Hebrew University, Givat Ram campus, by crossing positive males with C57BL/6OLAHSD female mice obtained from Harlan Laboratories. All animals (wild types and transgenic littermates of same sex) were group housed both before and during the experiments. They were maintained under standard environmental conditions-temperature (20–22°C), humidity (55 ± 10 %), and 12-12 h light/dark cycle (7am on and 7pm off), with ad libitum access to water and food. Behavioral assays were performed during the light phase of the circadian cycle. All animal protocols were approved by the Institutional Animal Care and Use Committees at the Hebrew University of Jerusalem and were in accordance with the National Institutes of Health Guide for the Care and Use of Laboratory Animals. Animals were randomly assigned to individual experimental groups, with some exceptions, such as in case of conditioned place preference experiments (elaborated later). Experimenters were blinded regarding experimental manipulations wherever possible.

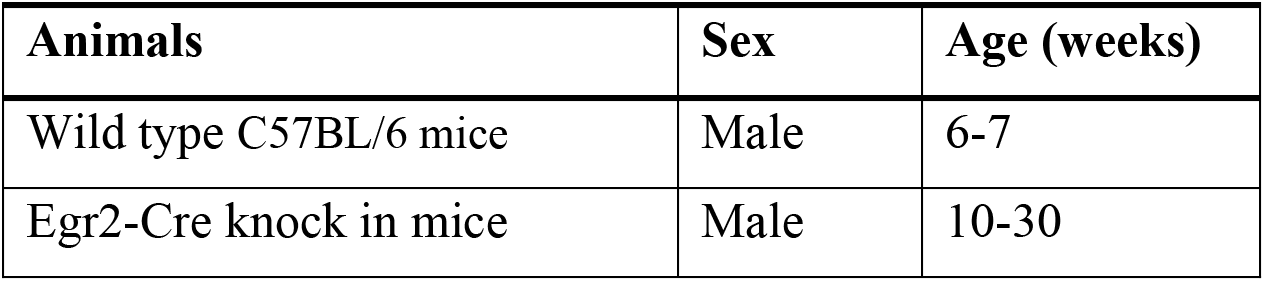

### Detailed methods

#### Behavioral Assays

##### Cocaine sensitization

6-7-week-old C57BL/6OLAHSD mice, after arriving from Harlan Laboratories were first allowed to acclimate to the SPF facility for a period of 5-7 days. Animals were then briefly handled once or twice daily for 2-3 days. During the handling sessions, animals were allowed to freely move around on the experimenter’s palm for 1-2 minutes either alone or in pairs. On the following 3 consecutive days, mice were subjected to once daily intraperitoneal (IP) saline injections (250 μl), and immediately transferred to a clear Plexiglas box (30×30×30 cm) within a sound- and light-attenuated chamber fitted with an overhead camera, for ~20 mins, and then returned to their home cage. After this habituation phase, animals were subjected to one daily IP cocaine injection (20 mg/kg; Stock solution: 2mg/ml dissolved in 0.9% saline and injected at 10ml/kg volume), according to the following groups: (1) *Acute cocaine* group received a single dose of IP cocaine, (2) *repeated cocaine* group were administered cocaine once daily for 5 consecutive days, and (3) *Challenge cocaine* group of animals were treated similarly to the repeated cocaine group for the first 5 days, subjected to abstinence (no drug treatment) for 21-22 days, and then re-exposed to a single dose of cocaine. Animals sacrificed directly from the home cage without any treatment were regarded as controls in the experiment (0h), and interleaved with the other groups corresponding to the relevant cocaine regiment (acute, chronic, and challenge cocaine). Transcription was analyzed at 1, 2, and 4 hours following the cocaine injection for the RNA-seq experiments. In smFISH experiments, animals were sacrificed for brain collection 1h after the cocaine injection, while control animals were treated as described earlier. Locomotor activity measured as distance travelled in the open field arena for a period of 15 mins, following either saline/cocaine injections, on each day was quantified by Ethovision (Noldus) software.

##### Conditioned Place Preference

Conditioned place preference was assessed in a custom-fitted arena [Plexiglass box (30×30×30 cm)] designed in-house, and placed in individual light- and sound-attenuated chambers [as in (Terem et al., 2020)]. On the preference test days, the arena was divided into two compartments of equal dimensions. One compartment was fitted with rough floor (“crushed ice” textured Plexiglas) and black (on white) dotted wallpaper, while the other was fitted with smooth floor with black (on white) striped wallpaper. On the conditioning days, animals were presented with only one context in each training session, such that the entire box had rough flooring and dotted wallpaper or smooth flooring with striped wallpaper. Animals were placed in the center of the arena and free behavior was recorded for 20 min. General activity and position/location of the mice in the arena was monitored by video recording using an overhead camera. Baseline preference was measured using the Ethovision XT software by analyzing the time spent in each chamber during the 20 mins session. Mice were randomly assigned a conditioning compartment in order to approximately balance any initial bias in preference towards a specific chamber. *Procedure:* All experiments were performed using an unbiased design, and consisted of the following phases: Handling: 2-3 days performed twice daily, and involved free exploration on the palms of the experimenter for 2-3 mins. Pre-test: Single 20 mins session (performed around noon), during which animals explored the arena which was divided into two compartments. Conditioning-3 days of 2 counterbalanced 20 min sessions per day separated by at least 4 hours. Mice were randomly assigned to a context (combination of a single floor-type and wallpaper patterns, as described above) which was paired with IP injections of saline (250 μl), and a separate context, which was paired with IP cocaine (10 mg/kg; Stock solution: 1mg/ml dissolved in 0.9% saline and injected at 10ml/kg volume). Post-conditioning final preference test was performed as in the Pre-test.

For chemogenetic experiments, Clozapine-N-Oxide (CNO; 10mg dissolved in 500μl DMSO and then mixed into 9.5 ml 0.9% saline, to a total of 10 ml CNO solution at a concentration of 1mg/ml. was injected at a 10mg/Kg dose 30 mins before cocaine conditioning sessions.

#### Tissue dissections and RNA extraction

Collection of tissue samples (Figure 3–figure supplement 1) and RNA-extraction were performed as described previously (Mukherjee et al., 2018; Turm et al., 2014) with few modifications. Briefly animals were anesthetized in isoflurane (Piramal Critical Care), euthanized by cervical dislocation, and the brains quickly transferred to ice cold artificial cerebrospinal fluid (ACSF) solution. Coronal slices of 400 μm were subsequently made on a vibrating microtome (7000 smz2; Camden Instruments) and relevant brain areas dissected under a stereoscope (Olympus). Tissue pieces were collected in PBS, snap-frozen in dry-ice and on the same day transferred to Tri-Reagent (Sigma-Aldrich). The tissue was stored at −80°C until being processed for RNA extraction. For RNA extraction, the stored tissue was thawed at 37°C using a drybath and then immediately homogenized using TissueLyser LT (Qiagen). RNA-extraction was performed according to the manufacturer’s guidelines. All steps were performed in cold conditions.

#### RNA-seq library preparation

100ng of RNA was used for first strand cDNA preparation as follows: The RNA was mixed with RT primers containing barcodes (7 bps) and unique molecular identifiers (UMIs; 8 bps) for subsequent de-multiplexing and correction for amplification biases, respectively. The mixture was denatured in a Thermocycler (Biorad) at 72°C for 3 minutes and transferred immediately to ice. An RT reaction cocktail containing 5X SmartScribe buffer, SmartScribe reverse transcriptase (Takara), 25mM dNTP mix (NEB), 100mM MnCl_2_ (Sigma) was added to the RNA and primer mix, and incubated at 42°C for 1 hour followed by 70°C for 15 minutes. The cDNA from all samples were pooled, cleaned with 1.2X AMPURE magnetic beads (Beckman Coulter) and eluted with 10mM Tris of pH-8 (ThermoFisher Scientific). The eluted cDNA was further processed for double-stranded DNA synthesis with the NEBNext Ultra II Non-Directional RNA Second Strand Synthesis Module (NEB), followed by another round of clean-up with 1.4X SPRI magnetic beads. The resultant double stranded cDNA was then incubated with Tn5 tagmentation enzyme and a 21bp oligo (TCGTCGGCAGCGTCAGATGTG sequence) at 55°C for 8 minutes. The reaction was stopped by denaturing the enzyme with 0.2% SDS (Biological Industries), followed by another round of cleaning with 2X SPRI magnetic beads. The elute was amplified using the KAPA Hifi Hotstart ReadyMix (Kapa Biosystems) and forward primer that contains Illumina P5-Read1 sequence) and reverse primer containing the P7-Read2 sequence. The resultant libraries were loaded on 4% agarose gel (Invitrogen) for size selection (250-700bp) and cleaned with Mini Elute Gel Extraction kit (Qiagen). Library concentration and molecular size were determined with NEBNext Library Quant Kit for Illumina (NEB) according to manufacturer’s guidelines, as well as Bioanalyzer using High Sensitivity DNA kit (Agilent Technologies). The libraries were run on the Illumina platform using NextSeq 500 High Output V2 kits (Illumina).

#### Single molecule fluorescence in-situ hybridization

A detailed protocol is available in (Gonzales et al., 2020). Briefly, smFISH protocol was performed on 14μm tissue sections using the RNAscope® Multiplex Fluorescent Reagent kit (Advanced Cell Diagnostics) according to the RNAscope® Sample Preparation and Pretreatment Guide for Fresh Frozen Tissue and the RNAscope® Fluorescent Multiplex Kit User Manual (Advanced Cell Diagnostics). Image acquisition was performed using a Hermes high definition cell-imaging system with 10x 0.4NA and 40X 0.75NA objectives. 5 Z-stack images were captured for each of 4 channels-475/28 nm (FITC), 549/15 nm (TRITC), 648/20 nm (Cy5) & 390/18 nm (DAPI). Image processing was performed using ImageJ software. Maximum intensity images for each channel were obtained using Maximum Intensity Z-projection. All channels were subsequently merged, and the dorsal striatum region was manually cropped from these merged images according to the Franklin and Paxinos Mouse brain atlas, Third edition. Quantification of RNA expression from images was done using the CellProfiler speckle counting pipeline.

#### Stereotactic surgeries

Induction and maintenance of anesthesia during surgery was achieved using SomnoSuite Low-Flow Anesthesia System (Kent Scientific Corporation). Following induction of anesthesia, animals were quickly secured to the stereotaxic apparatus (David KOPF instruments). The skin was cleaned with Betadine (Dr. Fischer Medical), and Lidocaine (Rafa Laboratories) was applied to minimize pain. An incision was made to expose the skull, which was immediately cleaned with Hydrogen peroxide (GADOT), and a small hole was drilled using a fine drill burr (RWD Life Science). Using a microsyringe (33G; Hamilton) connected to an UltraMicroPump (World Precision Instruments) virus was subsequently injected at a flow rate of 100nl/min. Upon completion of virus delivery, the microsyringe was left in the tissue for up to 5 minutes, and then slowly withdrawn. The skin incision was closed using a Vetbond bioadhesive (3M), the animals were removed from the stereotaxic apparatus, injected with saline and pain-killer Rimadyl (Norbrook), and allowed to recover under gentle heating. Coordinates of the stereotactic injection were determined using the Paxinos and Franklin mouse brain atlas. Every virus used in the study was titrated appropriately to ensure localized infections.

#### Coordinates of the stereotactic injection

**Coordinates of the stereotactic injection:**
| **Experiment ID** | **Viruses** | **Coordinates** | **Strain** |
| --- | --- | --- | --- |
| Kir2.1 inhibition<br>(**Fig 1I,J**) | AAVdj-DIO-eGFP (n= 3)<br>AAVdj-CAG-DIO-GPE2 (Kir2.1) (n= 3) | AP: 0.9; ML: $\pm 1.7$ & 2.4; DV: 3.2 & 3.5 | Egr2-Cre |
| Chemogenetic inhibition<br>(**Fig 3 D, E**) | AAV2-hSyn-DIO-hM4d(Gi)-mCherry (n= 8; all received CNO)<br>AAV2-hSyn-DIO- mCherry (n= 8, all received CNO) | AP: 0.9; ML: $\pm 2.6$; DV: 3.6 | Egr2-Cre |
| DN-Egr2 (**Fig 3F-H**) | AAVdj-CMV-eGFP (n= 8)<br>AAVdj-CAG-DNEgr2-IRES-GFP (n= 8) | AP: 0.9; ML: $\pm 2.65$; DV: 3.6 | WT |

### Quantification and Statistical Analysis

#### Statistical analysis and data visualization

R version 3.4.4 was used for all statistical analysis and graphical representations. Venn diagrams were generated with “eulerr” package. Three dimensional plots were generated with “plot3D” package. Heatmaps were generated with “Heatmap.2” function form “gplots” package. All other figures were generated using “ggplot2”. Details of the statistics applied in analysis of smFISH and behavioral experiments are summarized in Supplementary file 6.

#### RNA-seq analysis

##### Alignment and QC

RNAseq read quality was evaluated using FastQC. PCR duplicates were removed using Unique molecular identifiers (UMIs), and polyA tail, if existing, was trimmed from the 3’ end of the reads. Reads were aligned to the mouse genome (GRCm38) using STAR, and HTseq was used to count the number of reads for each gene. Samples with less than one million aligned reads were removed from the analysis. Samples with more than 8 million reads were down-sampled to 50% (using R package “subSeq”). The list of the samples analyzed in this paper and the distribution of library size are presented in Supplementary file 2 and Figure1–figure supplement 2. All raw sequencing data is available on NCBI GEO: GSE158588.

##### Analysis of shifts in baseline transcription

In order to compare baseline shifts in gene expression following repeated cocaine administration we compared gene expression within the samples obtained at time 0 (not exposed to cocaine on day of sample collection) in each one of the conditions – acute, repeated and challenge cocaine (Figure1–figure supplement 2B - heatmap of all genes exhibiting change). This analysis was performed with “DEseq2” package in R. We used the Wald test in the DEseq function and compared gene expression in cocaine naïve mice vs. mice exposed to repeated cocaine, as well as comparing to abstinent mice following repeated cocaine. List of detected genes, normalized counts and *p*-values (FDR corrected) are presented in Supplementary file 2. We observed that in a few samples an apparent sequencing batch effect was detected, likely related to the library preparation and/or to association of samples with different sequencing runs. Therefore, we performed the final analysis on only a subset of the samples, which did not exhibit a batch effect. While gene selection was performed on the subset of samples, the data portrayed in Figure 1C depicts all samples from the relevant time points – demonstrating that the genes identified from the subset of samples are consistently modified across all samples. Therefore, our gene list likely provides a conservative underestimate of the true magnitude of shifts in gene expression.

##### Analysis of inducible transcription

Detection of the induced genes following cocaine administration was performed with the “DEseq2” package in R. Each structure was analysed separately. The model included time (0,1,2,4 hours after cocaine administration) and the experiment (acute, repeated and challenge) as well as the interaction time x experiment. We used a likelihood ratio test (LRT) and selected genes changing over time in at least one of the experiments (eliminating genes that are changing only between experiments, but not in time). Next, to evaluate the effect of time in each specific experiment, we used the selected gene list and fitted a generalized linear model with a negative binomial distribution followed by LRT for each experiment separately. Genes with p<0.05 (corrected) and fold change > 1.2 were considered significant. List of the detected genes, normalized counts and *p*-values (FDR corrected) are presented in Supplementary file 4.

##### Gene annotation and functional analysis

KEGG pathway analysis was performed using the “SPIA” package (Signaling Pathway Impact Analysis) in R. Pathways with *p*<0.05 and at least eight differentially expressed genes were considered significant.

GO terms enrichment analysis was performed using the “clusterProfiler” package in R. Biological Process (BP) and Molecular Function (MF) sub-ontologies were included in the analysis. The results of the inducible transcription analysis (*p*<0.05, FDR corrected) are included in Supplementary file 5 (complete list of enriched GO terms and genes) and in Figure 1H (representative GO term list). In the analysis of baseline transcription, we perform a second step of clustering in order to remove redundancy and identify global patterns across structures. After selecting the significantly enriched GO terms (*p*<0.05, FDR corrected) we grouped together all GO terms that shared at least 50% identity of the differentially expressed genes in any of the structures (Supplementary file 3). As described in the results section, few clusters were selected for presentation, and the expression levels of genes included in these clusters - across all time points and all structures, are presented as a heatmap in Figure 1C and Figure1–figure supplement 4.

##### smFISH analysis

For the IEGs probes, selection for ‘robust-expressing’ cells was done as follows: We used the cocaine-naïve control data and after removing the non-expressing cells (cells expressing 0-1 puncta), the remaining cells were binned equally into three groups based on the per-cell expression levels, and the top 33% cells were defined ‘robust expressors’. Thus, cells qualified as ‘robust expressors’ for a given IEG if they expressed at least the following number of puncta per cell: *Arc* - 11, *Egr2* - 6, *Nr4a1*- 12, *Fos* - 5. For *Drd1* and *Drd2* expression, a threshold of 8 puncta/cell was implemented (Gonzales et al., 2020).

In order to identify the area with the highest density of IEG expressing cells in the striatum, we performed Two-Dimensional Kernel Density Estimation using the function ‘geom_density_2d’ in R [as in (Gonzales et al., 2020)]. This function estimates two-dimensional kernel density with an axis-aligned bivariate normal kernel, evaluated on a square grid, while displaying the result with contours. The regions of highest density, within which at least 20% of the cells are found, were selected. This process was performed independently for each one of the replicas and the selected contours plotted. A list of the samples and number of cells included in the analysis is found in Supplementary file 7. Details of statistical analysis and results for smFISH data are summarized in Supplementary file 6. Raw data (puncta per cell) is available on Mendeley Data (http://dx.doi.org/10.17632/p5tsv2wpmg.1)

## Acknowledgements

The authors appreciate the helpful critical comments of members of the Citri lab and Prof. Inbal Goshen on data, writing and presentation. Prof. Ido Amit generously provided instruction and guidance on RNAseq library preparation. Work in the Citri laboratory is funded by the European Research Council (ERC 770951), The Israel Science Foundation (1062/18, 393/12, 1796/12, and 2341/15), The Israel Anti-Drug Administration, EU Marie Curie (PCIG13-GA-2013-618201), the National Institute for Psychobiology in Israel, Hebrew University of Jerusalem Israel founded by the Charles E. Smith family (109-15-16), an Adelis Award for Advances in Neuroscience, the Brain and Behavior Foundation (NARSAD 18795), German–Israel Foundation (2299-2291.1/2011), and Binational Israel–United States Foundation (2011266), the Milton Rosenbaum Endowment Fund for Research in Psychiatry, a seed grant from the Eric Roland Fund for interdisciplinary research administered by the ELSC, contributions from anonymous philanthropists in Los Angeles and Mexico City, as well as research support from the Safra Center for Brain Sciences (ELSC) and the Canadian Institute for Advanced Research (CIFAR). DM was funded by a “Golden Opportunity Doctoral fellowship”, as well as a “Bridging” Post-Doctoral Fellowship from the Jerusalem Brain Community.

## Author contributions

Diptendu Mukherjee, Ben Jerry Gonzales & Reut Fluss, Conceptualization, Data curation, Formal analysis, Validation, Investigation, Visualization, Methodology, Writing - original draft, Writing - review and editing; Hagit Turm, Validation, Investigation, Methodology, Project administration; Maya Groysman, Resources, Methodology; Ami Citri, Conceptualization, Resources, Formal analysis, Supervision, Funding acquisition, Visualization, Methodology, Writing—original draft, Project administration, Writing—review and editing.

## Supplementary Figures

**Figure 1 – figure supplement 1:** Boundaries of dissected brain structures (related to Figure 1)

**Figure 1 – figure supplement 2:** Quality control analysis of RNA-seq experiments (related to Figure 1)

**Figure 1 – figure supplement 3:** Repeated cocaine administration and abstinence induce baseline shifts in gene transcription (related to Figure 1)

**Figure 1 – figure supplement 4:** Repeated cocaine exposure and abstinence alters the expression of gene clusters related to neuroplasticity (related to Figure 1)

**Figure 1 – figure supplement 5:** Domains of AAV transduction and expression in Egr2-CRE mice (related to Figure 1)

**Figure 2 – figure supplement 1:** Cocaine dynamically modulates cellular IEG expression in the VLS and MS (related to Figure 2)

**Figure 2 – figure supplement 2:** Coherence of cocaine-induced IEG expression (related to Figure 2)

**Figure 3– figure supplement 1:** Induction of IEGs are correlated to Drd1 expression in the VLS, and to both Drd1 & Drd2 expression in the MS (related to Figure 3)

**Figure 3 – figure supplement 2:** Drd1 and Drd2 receptor expression in the VLS and MS. (related to Figure 3)

**Figure 3– figure supplement 3**: DREADD inhibition of VLS Egr2^+^ cells does not affect locomotion. (related to Figure 3)

**Figure 3– figure supplement 4**: Disruption of Egr2 function in the VLS does not affect locomotor behavior. (related to Figure 3).

## Supplementary files

**Supplementary file 1:** Distribution of sequencing libraries analysed in this study

**Supplementary file 2:** Normalized reads of baseline shifted genes (tabs corresponding to individual structures)

**Supplementary file 3:** Clusters of Gene Ontology (GO) analysis and corresponding genes

**Supplementary file 4:** Normalized reads of induced genes (tabs correspond to structures)

**Supplementary file 5:** Gene Ontology annotation of cocaine-induced genes in the dorsal striatum

**Supplementary file 6:** Statistical analysis

**Supplementary file 7:** Distribution of cell numbers among replicates in smFISH analysis

---

**Figure 1 – figure supplement 1:**
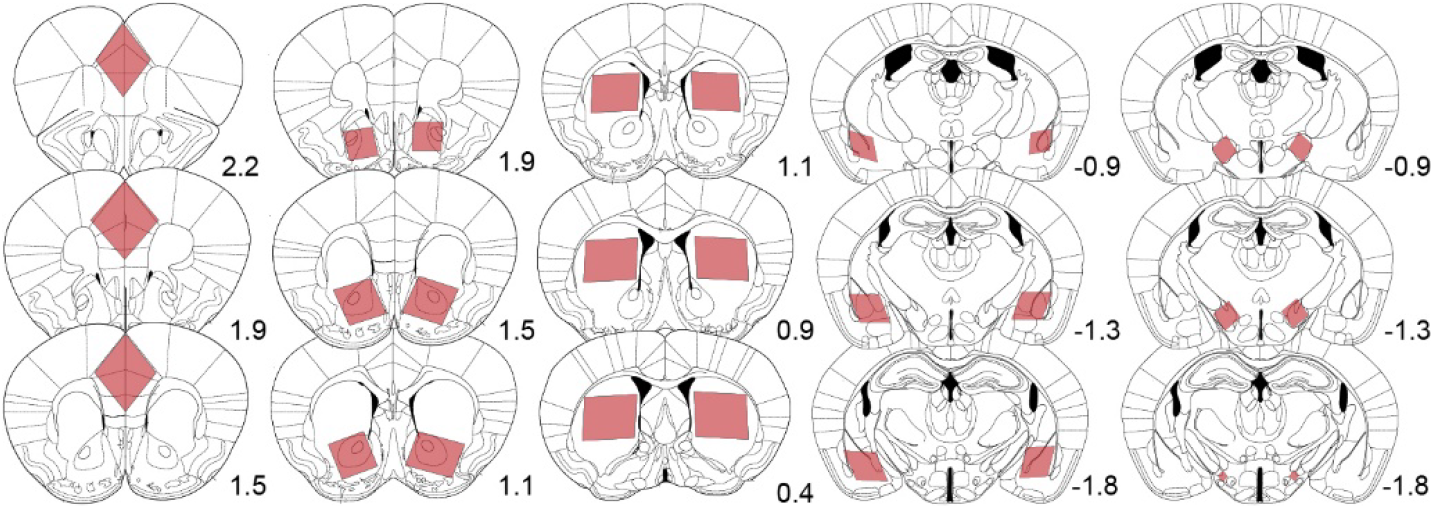
Boundaries of dissected brain structures. Illustration of the brain tissue collected for RNA extraction and sequencing analysis. Red shaded boxes represent the area dissected from 400-micron sections at specific distances from Bregma (marked by the number). Left to right, brain regions include limbic cortex (medial prefrontal and cingulate cortex together), nucleus accumbens, dorsal striatum, amygdala, and lateral hypothalamus.

**Figure 1 – figure supplement 2:**
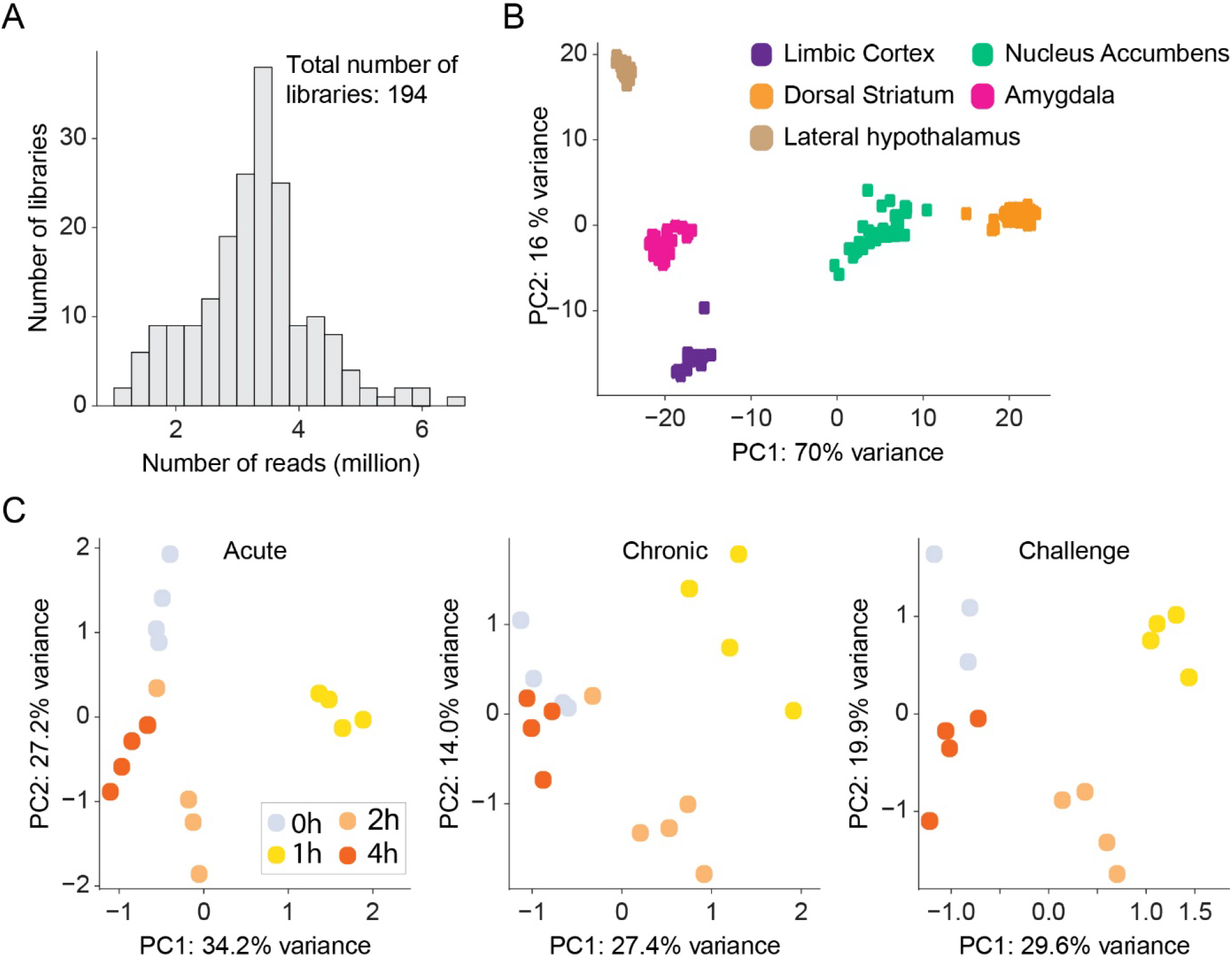
Quality control analysis of RNA-seq experiments. **(A)** Histograms of mean read depth of 3’-RNAseq libraries. n=194 libraries. **(B)** Principal component analysis (PCA) demonstrates clustering of RNAseq libraries according to brain region, consistent with brain nuclei expressing distinct transcriptional profiles (Hawrylycz et al., 2012; Kang et al., 2011; Ortiz et al., 2020). Each dot represents individual libraries color coded by brain structure. n=194 samples. **(C)** PCA of genes differentially expressed in the DS following acute, repeated and challenge cocaine reveals clustering according to the time point of gene expression. Libraries of 1h time-point are most distinct from the relevant control (0h) groups. Each dot represents a sample color coded according to the time point. n=48 samples. These analyses indicate the robustness of transcriptional dynamics induced by cocaine, and also provide an indication for the reproducibility of the experimental samples and the quality of the sequencing.

**Figure 1 – figure supplement 3:**
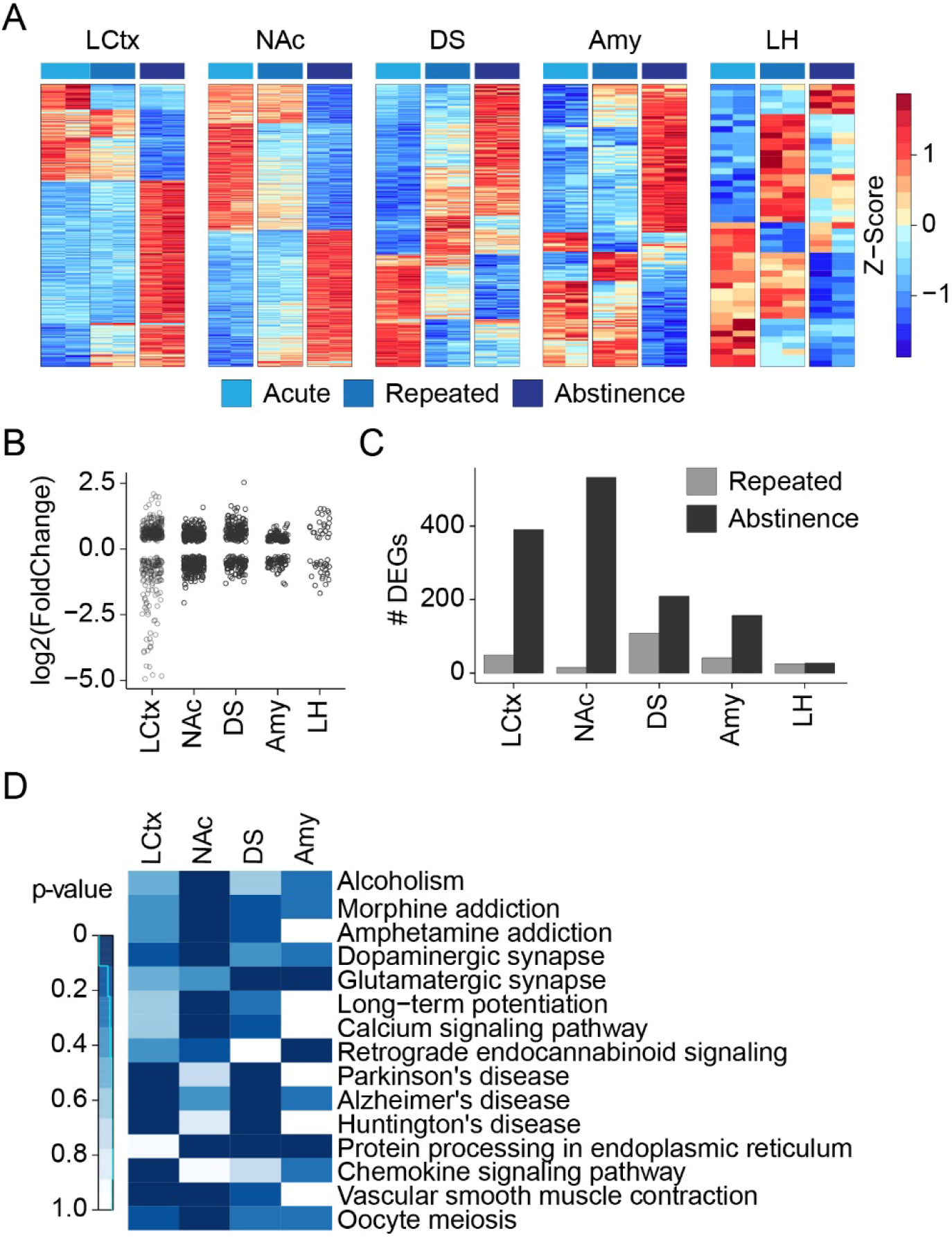
Repeated cocaine administration and abstinence induce baseline shifts in gene transcription. **(A)** Gene expression profiles of repeated cocaine treated and abstinent mice within different brain regions. Heatmaps reflect log2 (fold change) of genes demonstrating differential expression between 0h of acute (cocaine naïve), repeated and challenge cocaine in different brain regions. Columns represent individual samples sorted by the cocaine experience and brain nuclei. Expression is graded according to the color bar. Differentially-expressed genes (DEGs) were assayed from samples that were included in the same sequencing run in order to avoid the introduction of batch effects. n=2 samples in each condition. (FC>1.2; FDR corrected p<0.05 in at least one of the conditions, Wald test – see methods). LCtx: limbic cortex, NAc: nucleus accumbens, DS: dorsal striatum, Amy: amygdala, LH: lateral hypothalamus. **(B)** Dots-plots demonstrate the magnitude and direction of DEG expression shifts in different brain nuclei. Each dot represents a DEG. **(C)** The number of genes that exhibit shift in their baseline expression following repeated cocaine and abstinence. Bar graphs color coded by brain structure, depict the number of DEGs in each structure. **(D)** KEGG pathway annotation of gene clusters whose expression is modulated by repeated drug exposure and abstinence. Heat map showing enriched pathways consisting of at least 6 DEGs in different brain nuclei. Color represents p-value.

**Figure 1 – figure supplement 4:**
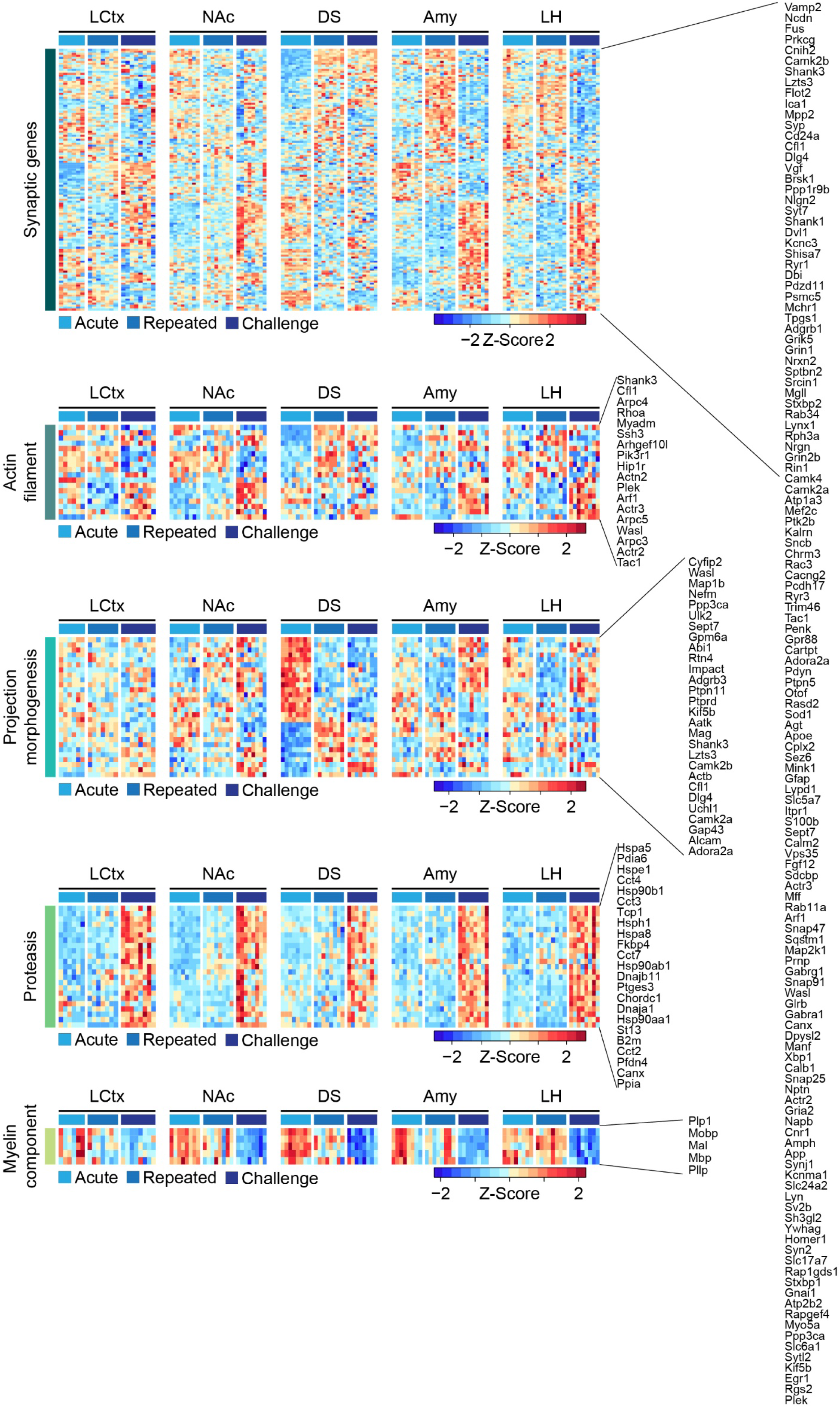
Repeated cocaine exposure and abstinence alters the expression of gene clusters related to neuroplasticity. Heatmaps depict the expression levels of genes which are modulated between naïve mice and mice following repeated cocaine or abstinence in different brain regions clustered according to GO terms related with neuroplasticity (from Fig. 1C). Columns represent individual mice, while expression is graded according to the color bar (Z-scored per gene). Cocaine experiences are color coded according to the key.

**Figure 1 – figure supplement 5:**
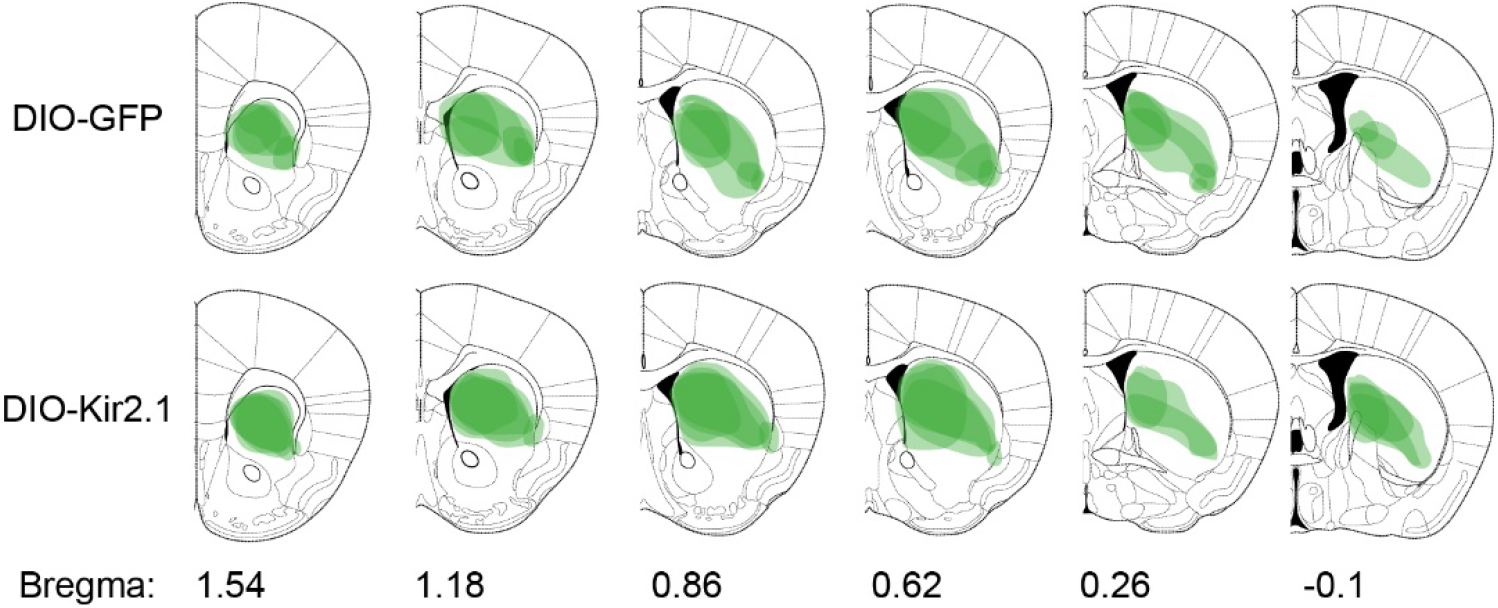
Domains of AAV transduction and expression in Egr2-CRE mice. Summary of AAV-DIO-GFP and AAV-DIO-Kir2.1 transduction domains in the striatum of Egr2-Cre mice. Outlines of the infections were manually overlaid on equivalent coronal sections from the mouse brain atlas. n= 3 mice in each group.

**Figure 2 – figure supplement 1:**
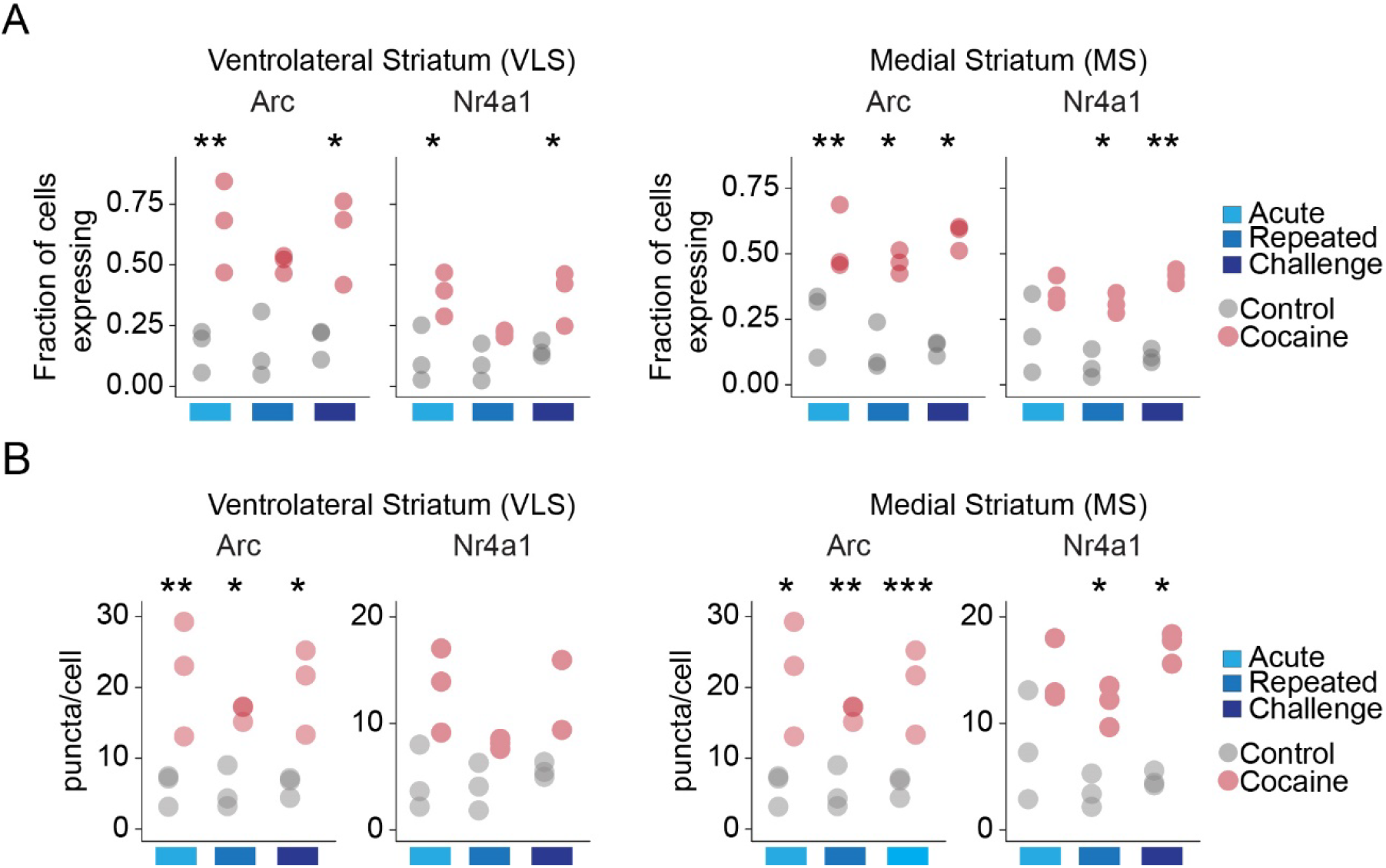
Cocaine dynamically modulates cellular IEG expression in the VLS and MS. (**A**) Dot-plot of the fraction of cells positive for expression of *Arc* and *Nr4a1* in the VLS and MS (threshold: *Arc*=11, *Nr4a1*=12 puncta/cell) following acute, repeated and challenge cocaine exposures. n=3 sections from 3 mice (8823-12246 cells). ANOVA with post hoc Tukey; \**p*<0.05, \*\**p*<0.005. (**B**) Dot-plot of the puncta/cell for *Arc* and *Nr4a1* following acute, repeated and challenge cocaine exposures. n=3 sections from 3 mice (8823-12246 cells). ANOVA with post hoc Tukey; \**p*<0.05, \*\**p*<0.005, *\*\*\*p<*0.005.

**Figure 2 – figure supplement 2:**
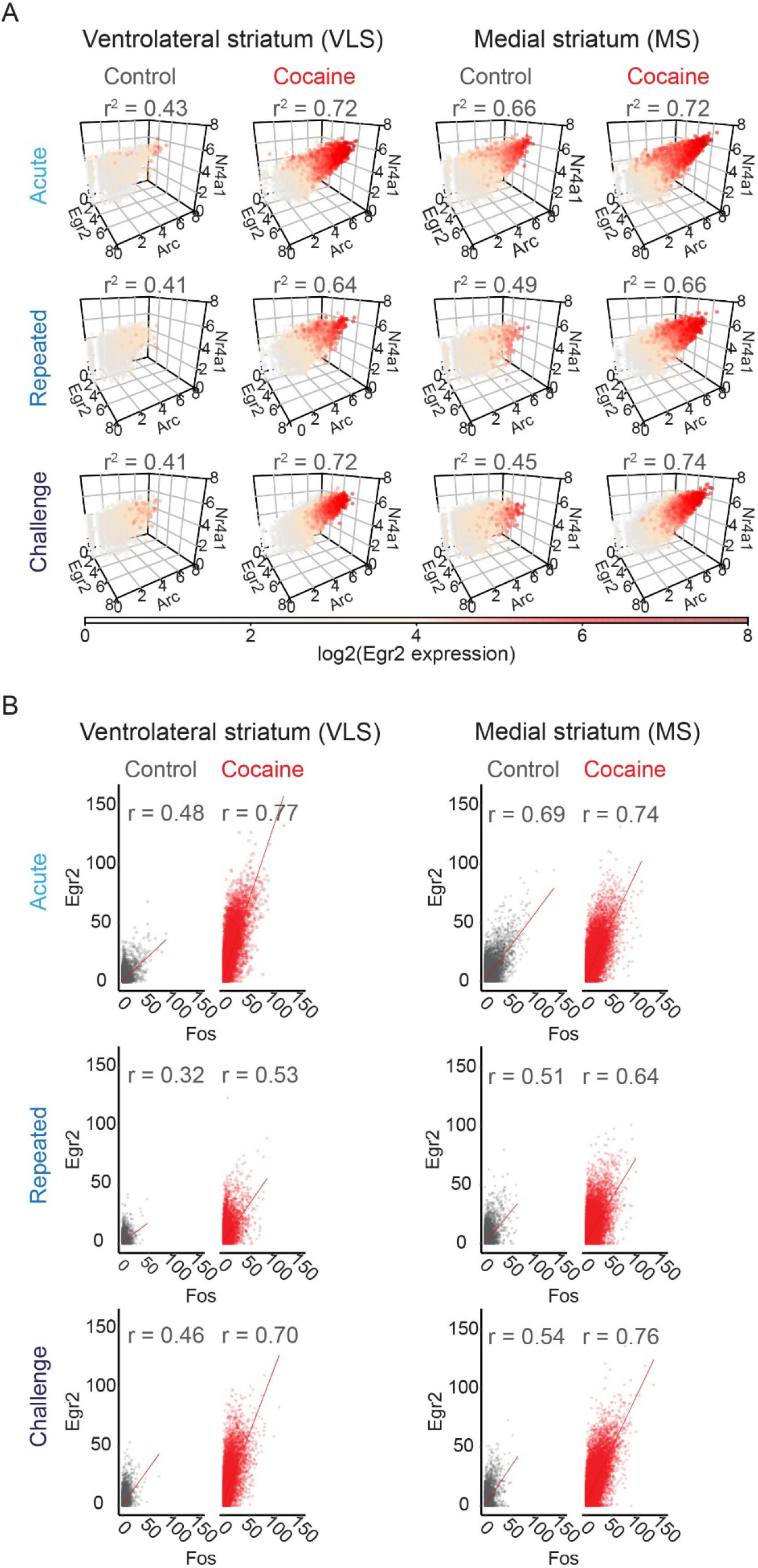
Coherence of cocaine-induced IEG expression. **(A)** *Arc*, *Egr2*, and *Nr4a1* are coherently induced in cells of the VLS and MS. Coherence refers to the correlated co-expression of multiple IEGs within individual cells. Each dot depicts the expression [log2 (puncta per cell + 1)] of *Arc*, *Egr2*, and *Nr4a1* within a single cell. The plots are color coded by *Egr2* expression according to the color bar. Data summarized from n=3 animals in each condition. Analysis using a linear model with *Egr2* expression as the dependent variable and *Arc, Nr4a1* expression as the independent variables demonstrated significant correlation between the genes (p<0.0001 for all conditions; 6850-16952 cells). **(B)** *Egr2* and *Fos* expression are strongly and positively correlated in the VLS and MS. Individual dots depict pairwise correlations of *Egr2* and *Fos* within single cells. n = 6 sections from 3 mice in each condition (Pearson correlation, p < 0.0001 for all conditions; 17734 - 33195 cells).

**Figure 3 – figure supplement 1:**
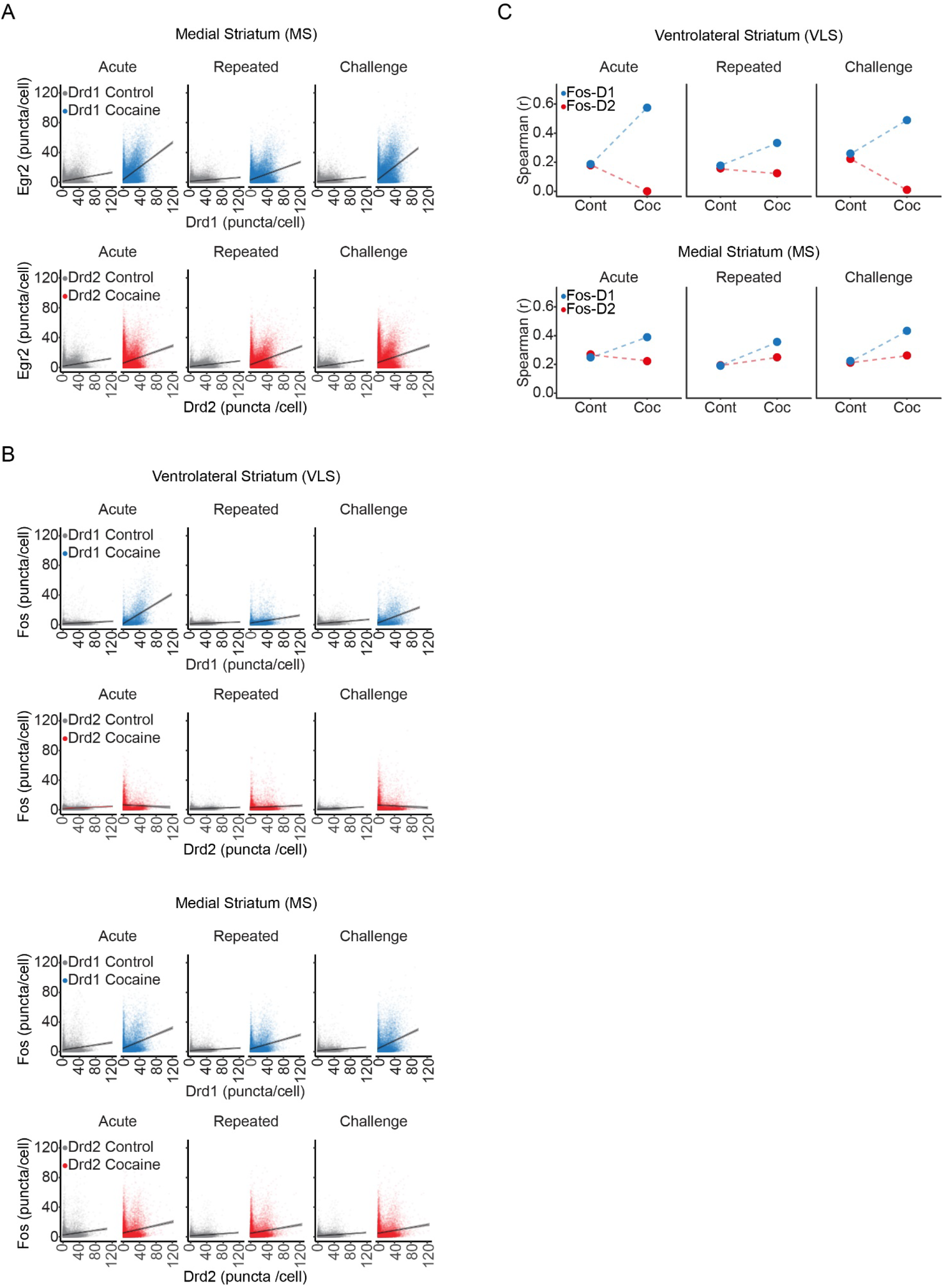
Induced IEGs are correlated to *Drd1* expression in the VLS, and to both *Drd1* & *Drd2* expression in the MS. **(A)** *Egr2* induction does not show cell-type bias in the MS, and is unaffected by the history of cocaine experience. Scatter plots show cellular *Egr2* expression with *Drd1* or *Drd2* expression (puncta/cell) within individual cells. n=6 sections from 3 mice for each condition (grey dots represent cells from 0h for either *Drd1* or *Drd2* combination, and blue or red dots for *Drd1* or *Drd2* combination, respectively, 1h following cocaine experience). Refer to Supplementary file 6 for detailed statistics. **(B)** Scatter plots of cellular expression (puncta/cell) of *Fos* with *Drd1* or *Drd2* in the VLS and MS. VLS-Fos induction following acute and challenge cocaine is biased to Drd1+ neurons. In the MS, irrespective of the cocaine experience, *Fos* induction occurs in both Drd1+ and Drd2+ neurons. Each dot represents a cell color coded similar to A. n=3 sections from 3 mice for each condition. **(C)** Line graphs depicting spearman correlation show specific and dramatic improvement of *Egr2* correlation with *Drd1* and not *Drd2* expression upon acute cocaine exposure in the VLS, which is dampened following repeated exposure and re-emerges upon cocaine challenge. In the MS, *Egr2* expression is consistently correlated to both *Drd1* and *Drd2* expression irrespective of the cocaine experience, with a modest bias towards Drd1+ neurons.

**Figure 3 – figure supplement 2:**
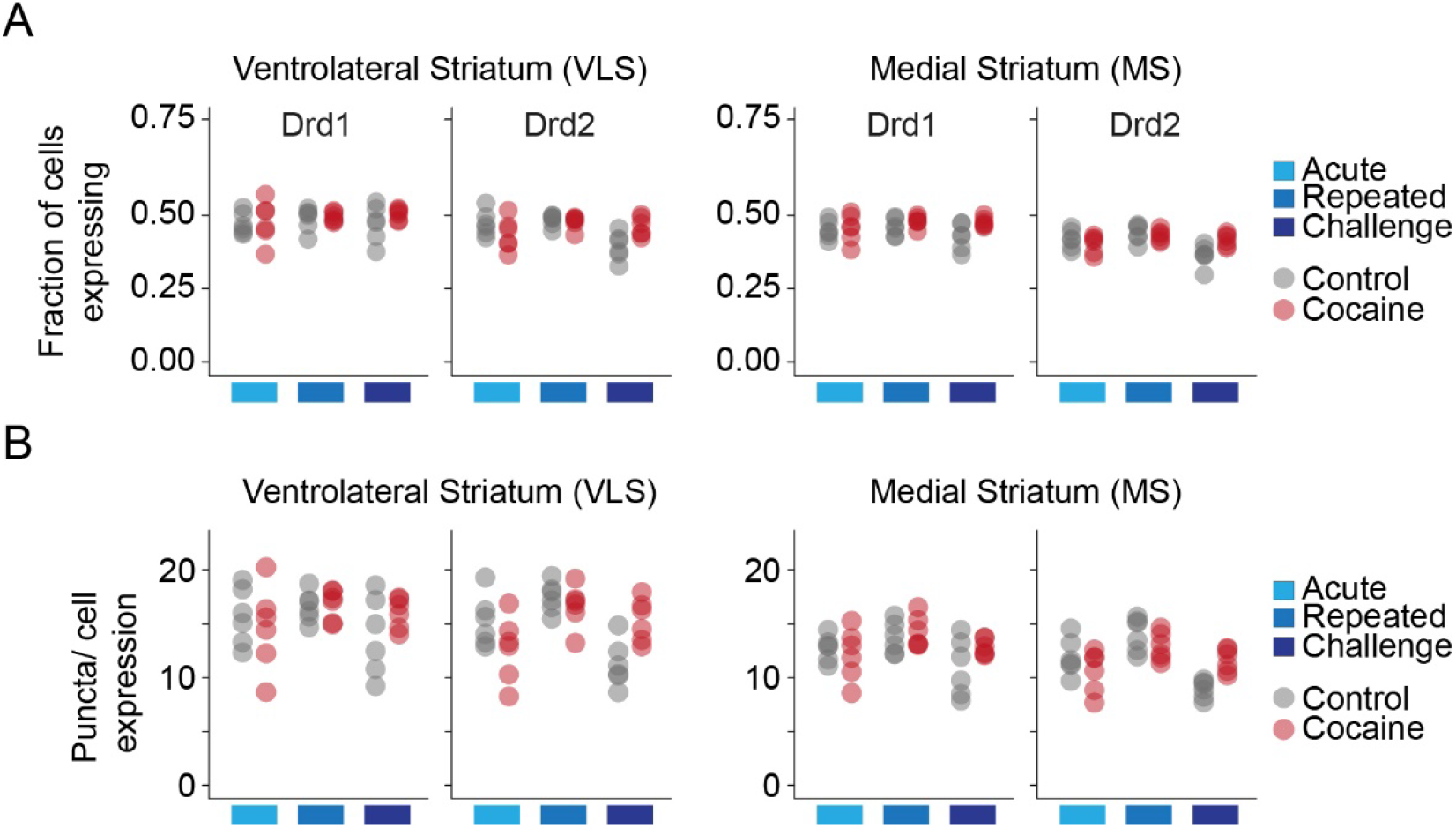
*Drd1* and *Drd2* receptor expression in the VLS and MS. **(A)** Dot-plots depicting the fraction of *Drd1* vs *Drd2* expressing cells in the VLS and MS in control (grey) and cocaine (red) conditions following distinct cocaine experiences. n = 6 sections from 3 mice in each condition. **(B)** Dot plots depicting puncta/cell expression of *Drd1* and *Drd2* following distinct cocaine experiences. Color code is similar to A. n = 6 sections from 3 mice in each condition. Data analyzed in A and B using Student’s t-test, See Supplementary file 6 for details.

**Figure 3 – figure supplement 3:**
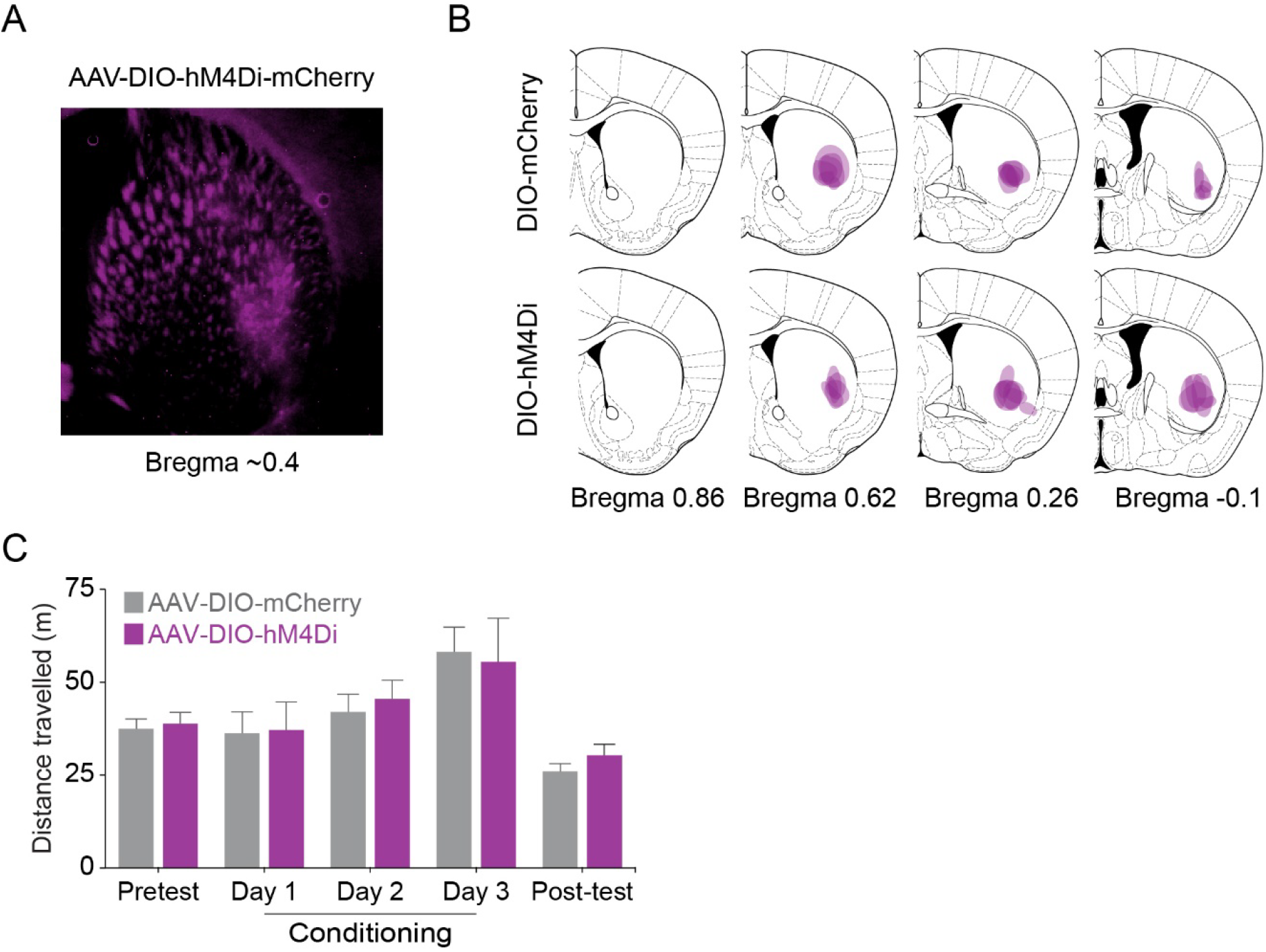
DREADD inhibition of VLS Egr2^+^ cells does not affect locomotion. **(A, B)** Verification of AAV transduction in the VLS of transgenic Egr2-CRE mice. Example of AAV-DIO-hM4Di-mcherry infections in the VLS (**A**) and summary of AAV-DIO-mCherry and AAV-DIO-hM4Di-mcherry infections (**B**). Span of viral transduction in individual mice was manually overlaid on corresponding coronal sections from the mouse brain atlas. n= 7 mice in each group. **(C)** Expression of AAV-DIO-hM4Di does not affect locomotor behavior (p = 0.93, ANOVA for interaction day with group, see Supplementary file 6 for stats). Bar graphs compare the locomotion of VLS-Egr2^mCherry^ and VLS-Egr2^hM4Di^ mice in the CPP chambers on pretest, conditioning session with cocaine, and post-test days. CNO (10 mg/kg, i.p.) was administered to both groups 30 min prior to cocaine conditioning. n= 7 mice in each group. Data represented as mean ± sem.

**Figure 3 – figure supplement 4:**
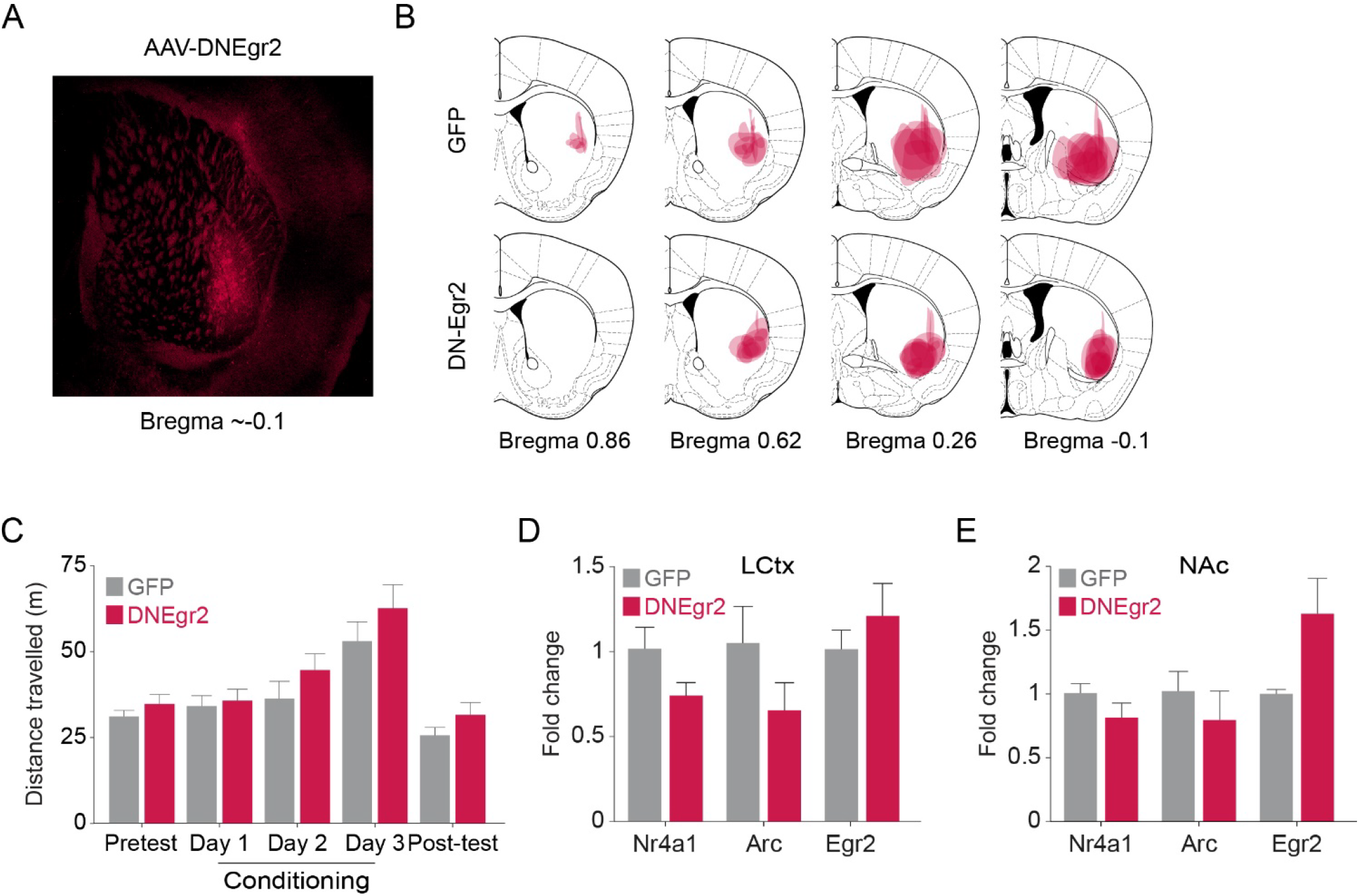
Disruption of Egr2 function in the VLS does not affect locomotor behavior. **(A, B)** Verification of AAV-GFP and AAV-DNEgr2 infection in the VLS. Example of AAV-DNEgr2 infection in the VLS **(A)**. **(B)** Summary of infection from individual mice of VLS^DNEgr2^ and VLS^GFP^ groups manually overlaid onto corresponding coronal sections from the mouse brain atlas. n= 8 mice in each group. **(C)** DN-Egr2 expression does not affect locomotor behavior (p = 0.7, ANOVA for interaction day with group, see Supplementary file 6). Bar graphs compare the average locomotion of VLS^GFP^ and VLS^DNEgr2^ mice in the CPP chambers on pretest, conditioning session with cocaine, and post-test days. n= 8 mice in each group. Data represented as mean ± sem. **(D)** Viral transduction of AAV-DNEgr2 in the VLS does not impact gene transcription in LCtx and NAc. Bar graphs represent expression of *Arc*, *Nr4a1* and *Egr2* in LCtx and NAc of VLS^GFP^ (grey) and VLS^DNEgr2^ (maroon) mice. n= 3 mice in each group. Data represented as mean ± sem.

## References

Alberini CM. 2009. Transcription Factors in Long-Term Memory and Synaptic Plasticity. Physiol Rev 89:121–145. doi:10.1152/physrev.00017.2008

Alberini CM, Kandel ER. 2015. The Regulation of Transcription in Memory Consolidation. Cold Spring Harb Perspect Biol 7:a021741. doi:10.1101/cshperspect.a021741

Albertson DN, Pruetz B, Schmidt CJ, Kuhn DM, Kapatos G, Bannon MJ. 2004. Gene expression profile of the nucleus accumbens of human cocaine abusers: evidence for dysregulation of myelin. J Neurochem 88:1211–1219. doi:10.1046/j.1471-4159.2003.02247.x

Amit I, Citri A, Shay T, Lu Y, Katz M, Zhang F, Tarcic G, Siwak D, Lahad J, Jacob-Hirsch J, Amariglio N, Vaisman N, Segal E, Rechavi G, Alon U, Mills GB, Domany E, Yarden Y. 2007. A module of negative feedback regulators defines growth factor signaling. Nat Genet 39:503–512. doi:10.1038/ng1987

Atlan G, Terem A, Peretz-Rivlin N, Sehrawat K, Gonzales BJ, Pozner G, Tasaka G, Goll Y, Refaeli R, Zviran O, Lim BK, Groysman M, Goshen I, Mizrahi A, Nelken I, Citri A. 2018. The Claustrum Supports Resilience to Distraction. Curr Biol 28:2752-2762.e7. doi:10.1016/j.cub.2018.06.068

Baker DA, Specio SE, Tran-Nguyen LTL, Neisewander JL. 1998. Amphetamine Infused Into the Ventrolateral Striatum Produces Oral Stereotypies and Conditioned Place Preference. Pharmacol Biochem Behav 61:107–111. doi:10.1016/S0091-3057(98)00070-7

Balleine BW, O’Doherty JP. 2010. Human and Rodent Homologies in Action Control: Corticostriatal Determinants of Goal-Directed and Habitual Action. Neuropsychopharmacology 35:48–69. doi:10.1038/npp.2009.131

Balleine BW, Ostlund SB. 2007. Still at the Choice-Point: Action Selection and Initiation in Instrumental Conditioning. Ann N Y Acad Sci 1104:147–171. doi:10.1196/annals.1390.006

Bannon M, Kapatos G, Albertson D. 2005. Gene Expression Profiling in the Brains of Human Cocaine Abusers. Addict Biol 10:119–126. doi:10.1080/13556210412331308921

Bannon MJ, Johnson MM, Michelhaugh SK, Hartley ZJ, Halter SD, David JA, Kapatos G, Schmidt CJ. 2014. A Molecular Profile of Cocaine Abuse Includes the Differential Expression of Genes that Regulate Transcription, Chromatin, and Dopamine Cell Phenotype. Neuropsychopharmacology 39:2191–2199. doi:10.1038/npp.2014.70

Bariselli S, Fobbs WC, Creed MC, Kravitz AV. 2019. A competitive model for striatal action selection. Brain Res 1713:70–79. doi:10.1016/j.brainres.2018.10.009

Bobadilla A-C, Dereschewitz E, Vaccaro L, Heinsbroek JA, Scofield MD, Kalivas PW. 2020. Cocaine and sucrose rewards recruit different seeking ensembles in the nucleus accumbens core. Mol Psychiatry. doi:10.1038/s41380-020-00888-z

Boerkoel CF, Takashima H, Bacino CA, Daentl D, Lupski JR. 2001. EGR2 mutation R359W causes a spectrum of Dejerine-Sottas neuropathy. Neurogenetics 3:153–157. doi:10.1007/s100480100107

Calipari ES, Bagot RC, Purushothaman I, Davidson TJ, Yorgason JT, Peña CJ, Walker DM, Pirpinias ST, Guise KG, Ramakrishnan C, Deisseroth K, Nestler EJ. 2016. In vivo imaging identifies temporal signature of D1 and D2 medium spiny neurons in cocaine reward. Proc Natl Acad Sci 113:2726–2731. doi:10.1073/pnas.1521238113

Caprioli D, Venniro M, Zhang M, Bossert JM, Warren BL, Hope BT, Shaham Y. 2017. Role of Dorsomedial Striatum Neuronal Ensembles in Incubation of Methamphetamine Craving after Voluntary Abstinence. J Neurosci 37:1014–1027. doi:10.1523/JNEUROSCI.3091-16.2017

Caster JM, Kuhn CM. 2009. Maturation of coordinated immediate early gene expression by cocaine during adolescence. Neuroscience 160:13–31. doi:10.1016/j.neuroscience.2009.01.001

Chandra R, Francis TC, Konkalmatt P, Amgalan A, Gancarz AM, Dietz DM, Lobo MK. 2015. Opposing Role for Egr3 in Nucleus Accumbens Cell Subtypes in Cocaine Action. J Neurosci 35:7927–7937. doi:10.1523/JNEUROSCI.0548-15.2015

Chandra R, Lobo MK. 2017. Beyond Neuronal Activity Markers: Select Immediate Early Genes in Striatal Neuron Subtypes Functionally Mediate Psychostimulant Addiction. Front Behav Neurosci 11:1–6. doi:10.3389/fnbeh.2017.00112

Clayton DF. 2000. The Genomic Action Potential. Neurobiol Learn Mem 74:185–216. doi:10.1006/nlme.2000.3967

Clayton DF, Anreiter I, Aristizabal M, Frankland PW, Binder EB, Citri A. 2020. The role of the genome in experience-dependent plasticity: Extending the analogy of the genomic action potential. Proc Natl Acad Sci 117:23252–23260. doi:10.1073/pnas.1820837116

Crombag H. 2002. Effect of Dopamine Receptor Antagonists on Renewal of Cocaine Seeking by Reexposure to Drug-associated Contextual Cues. Neuropsychopharmacology 27:1006–1015. doi:10.1016/S0893-133X(02)00356-1

Crombag HS, Bossert JM, Koya E, Shaham Y. 2008. Context-induced relapse to drug seeking: a review. Philos Trans R Soc B Biol Sci 363:3233–3243. doi:10.1098/rstb.2008.0090

Crombag HS, Shaham Y. 2002. Renewal of drug seeking by contextual cues after prolonged extinction in rats. Behav Neurosci 116:169–173. doi:10.1037/0735-7044.116.1.169

Cruz FC, Babin KR, Leao RM, Goldart EM, Bossert JM, Shaham Y, Hope BT. 2014. Role of Nucleus Accumbens Shell Neuronal Ensembles in Context-Induced Reinstatement of Cocaine-Seeking. J Neurosci 34:7437–7446. doi:10.1523/JNEUROSCI.0238-14.2014

Cruz FC, Javier Rubio F, Hope BT. 2015. Using c-fos to study neuronal ensembles in corticostriatal circuitry of addiction. Brain Res 1628:157–173. doi:10.1016/j.brainres.2014.11.005

Cruz FC, Koya E, Guez-Barber DH, Bossert JM, Lupica CR, Shaham Y, Hope BT. 2013. New technologies for examining the role of neuronal ensembles in drug addiction and fear. Nat Rev Neurosci 14:743–754. doi:10.1038/nrn3597

De S, Turman, Jr. JE. 2005. Krox-20 gene expression: Influencing hindbrain-craniofacial developmental interactions. Arch Histol Cytol 68:227–234. doi:10.1679/aohc.68.227

Delfs JM, Kelley AE. 1990. The role of D1 and D2 dopamine receptors in oral stereotypy induced by dopaminergic stimulation of the ventrolateral striatum. Neuroscience 39:59–67. doi:10.1016/0306-4522(90)90221-O

DeNardo L, Luo L. 2017. Genetic strategies to access activated neurons. Curr Opin Neurobiol 45:121–129. doi:10.1016/j.conb.2017.05.014

Dong Y, Nestler EJ. 2014. The neural rejuvenation hypothesis of cocaine addiction. Trends Pharmacol Sci 35:374–383. doi:10.1016/j.tips.2014.05.005

Eipper-Mains JE, Kiraly DD, Duff MO, Horowitz MJ, McManus CJ, Eipper BA, Graveley BR, Mains RE. 2013. Effects of cocaine and withdrawal on the mouse nucleus accumbens transcriptome. Genes, Brain Behav 12:21–33. doi:10.1111/j.1601-183X.2012.00873.x

Everitt BJ. 2014. Neural and psychological mechanisms underlying compulsive drug seeking habits and drug memories – indications for novel treatments of addiction. Eur J Neurosci 40:2163–2182. doi:10.1111/ejn.12644

Freeman WM, Lull ME, Patel KM, Brucklacher RM, Morgan D, Roberts DCS, Vrana KE. 2010. Gene expression changes in the medial prefrontal cortex and nucleus accumbens following abstinence from cocaine self-administration. BMC Neurosci 11:29. doi:10.1186/1471-2202-11-29

Gao P, de Munck JC, Limpens JHWW, Vanderschuren LJMJMJ, Voorn P. 2017a. A neuronal activation correlate in striatum and prefrontal cortex of prolonged cocaine intake. Brain Struct Funct 222:3453–3475. doi:10.1007/s00429-017-1412-4

Gao P, Limpens JHW, Spijker S, Vanderschuren LJMJ, Voorn P. 2017b. Stable immediate early gene expression patterns in medial prefrontal cortex and striatum after long-term cocaine self-administration. Addict Biol 22:354–368. doi:10.1111/adb.12330

García-Fuster MJ, Flagel SB, Mahmood ST, Watson SJ, Akil H. 2012. Cocaine Withdrawal Causes Delayed Dysregulation of Stress Genes in the Hippocampus. PLoS One 7:e42092. doi:10.1371/journal.pone.0042092

Gass P, Herdegen T, Bravo R, Kiessling M. 1994. High induction threshold for transcription factor KROX-20 in the rat brain: partial co-expression with heat shock protein 70 following limbic seizures. Mol Brain Res 23:292–298. doi:10.1016/0169-328X(94)90238-0

Gipson CD, Kupchik YM, Shen H, Reissner KJ, Thomas CA, Kalivas PW. 2013. Relapse Induced by Cues Predicting Cocaine Depends on Rapid, Transient Synaptic Potentiation. Neuron 77:867–872. doi:10.1016/j.neuron.2013.01.005

Gonzales BJ, Mukherjee D, Ashwal-Fluss R, Loewenstein Y, Citri A. 2020. Subregion-specific rules govern the distribution of neuronal immediate-early gene induction. Proc Natl Acad Sci 117:23304–23310. doi:10.1073/pnas.1913658116

Gray JM, Spiegel I. 2019. Cell-type-specific programs for activity-regulated gene expression. Curr Opin Neurobiol 56:33–39. doi:10.1016/j.conb.2018.11.001

Gremel CM, Lovinger DM. 2017. Associative and sensorimotor cortico-basal ganglia circuit roles in effects of abused drugs. Genes, Brain Behav 16:71–85. doi:10.1111/gbb.12309

Guez-Barber D, Fanous S, Golden SA, Schrama R, Koya E, Stern AL, Bossert JM, Harvey BK, Picciotto MR, Hope BT. 2011. FACS Identifies Unique Cocaine-Induced Gene Regulation in Selectively Activated Adult Striatal Neurons. J Neurosci 31:4251–4259. doi:10.1523/JNEUROSCI.6195-10.2011

Han M-H, Russo SJ, Nestler EJ. 2019. Molecular, Cellular, and Circuit Basis of Depression Susceptibility and Resilience. Neurobiology of Depression. Elsevier. pp. 123–136. doi:10.1016/B978-0-12-813333-0.00012-3

Hope BT. 2019. Fos-Expressing Neuronal Ensembles in Addiction Research. Neural Mechanisms of Addiction. Elsevier. pp. 75–88. doi:10.1016/B978-0-12-812202-0.00006-3

Hope BT, Nye HE, Kelz MB, Self DW, Iadarola MJ, Nakabeppu Y, Duman RS, Nestler EJ. 1994. Induction of a long-lasting AP-1 complex composed of altered Fos-like proteins in brain by chronic cocaine and other chronic treatments. Neuron 13:1235–1244. doi:10.1016/0896-6273(94)90061-2

Hrvatin S, Hochbaum DR, Nagy MA, Cicconet M, Robertson K, Cheadle L, Zilionis R, Ratner A, Borges-Monroy R, Klein AM, Sabatini BL, Greenberg ME. 2018. Single-cell analysis of experience-dependent transcriptomic states in the mouse visual cortex. Nat Neurosci 21:120–129. doi:10.1038/s41593-017-0029-5

Hurd YL, Herkenham M. 1993. Molecular alterations in the neostriatum of human cocaine addicts. Synapse 13:357–369. doi:10.1002/syn.890130408

Hyman SE. 2005. Addiction: A Disease of Learning and Memory. Am J Psychiatry 162:1414–1422. doi:10.1176/appi.ajp.162.8.1414

Hyman SE, Malenka RC, Nestler EJ. 2006. Neural Mechanisms of Addiction: The Role of Reward-Related Learning and Memory. Annu Rev Neurosci 29:565–598. doi:10.1146/annurev.neuro.29.051605.113009

Imperio CG, McFalls AJ, Hadad N, Blanco-Berdugo L, Masser DR, Colechio EM, Coffey AA, Bixler G V., Stanford DR, Vrana KE, Grigson PS, Freeman WM. 2018. Exposure to environmental enrichment attenuates addiction-like behavior and alters molecular effects of heroin self-administration in rats. Neuropharmacology 139:26–40. doi:10.1016/j.neuropharm.2018.06.037

Johnson MM, David JA, Michelhaugh SK, Schmidt CJ, Bannon MJ. 2012. Increased Heat Shock Protein 70 Gene Expression in the Brains of Cocaine-Related Fatalities may be Reflective of Postdrug Survival and Intervention rather than Excited Delirium. J Forensic Sci 57:1519–1523. doi:10.1111/j.1556-4029.2012.02212.x

Karler R, Calder LD, Thai LH, Bedingfield JB. 1994. A dopaminergic-glutamatergic basis for the action of amphetamine and cocaine. Brain Res 658:8–14. doi:10.1016/S0006-8993(09)90003-8

Kelley AE. 2004. Memory and Addiction. Neuron 44:161–179. doi:10.1016/j.neuron.2004.09.016

Kelley AE, Delfs JM. 1991. Dopamine and conditioned reinforcement. Psychopharmacology (Berl) 103:187–196. doi:10.1007/BF02244202

Kovalevich J, Corley G, Yen W, Rawls SM, Langford D. 2012. Cocaine-Induced Loss of White Matter Proteins in the Adult Mouse Nucleus Accumbens Is Attenuated by Administration of a β-Lactam Antibiotic during Cocaine Withdrawal. Am J Pathol 181:1921–1927. doi:10.1016/j.ajpath.2012.08.013

Koya E, Golden SA, Harvey BK, Guez-Barber DH, Berkow A, Simmons DE, Bossert JM, Nair SG, Uejima JL, Marin MT, Mitchell TB, Farquhar D, Ghosh SC, Mattson BJ, Hope BT. 2009. Targeted disruption of cocaine-activated nucleus accumbens neurons prevents context-specific sensitization. Nat Neurosci 12:1069–1073. doi:10.1038/nn.2364

Kravitz A V, Freeze BS, Parker PRL, Kay K, Thwin MT, Deisseroth K, Kreitzer AC. 2010. Regulation of parkinsonian motor behaviours by optogenetic control of basal ganglia circuitry. Nature 466:622–626. doi:10.1038/nature09159

Kreitzer AC, Malenka RC. 2008. Striatal Plasticity and Basal Ganglia Circuit Function. Neuron 60:543–554. doi:10.1016/j.neuron.2008.11.005

Kyrke-Smith M, Williams JM. 2018. Bridging Synaptic and Epigenetic Maintenance Mechanisms of the Engram. Front Mol Neurosci 11. doi:10.3389/fnmol.2018.00369

LeBlanc SE, Ward RM, Svaren J. 2007. Neuropathy-Associated Egr2 Mutants Disrupt Cooperative Activation of Myelin Protein Zero by Egr2 and Sox10. Mol Cell Biol 27:3521–3529. doi:10.1128/MCB.01689-06

Lee JLC. 2006. Cue-Induced Cocaine Seeking and Relapse Are Reduced by Disruption of Drug Memory Reconsolidation. J Neurosci 26:5881–5887. doi:10.1523/JNEUROSCI.0323-06.2006

Li K, Wu Y, Li Y, Yu Q, Tian Z, Wei H, Qu K. 2019. Landscape and dynamics of the transcriptional regulatory network during natural killer cell differentiation. bioRxiv. doi:10.1101/572768

Li X, Rubio FJ, Zeric T, Bossert JM, Kambhampati S, Cates HM, Kennedy PJ, Liu Q-R, Cimbro R, Hope BT, Nestler EJ, Shaham Y. 2015. Incubation of Methamphetamine Craving Is Associated with Selective Increases in Expression of Bdnf and Trkb, Glutamate Receptors, and Epigenetic Enzymes in Cue-Activated Fos-Expressing Dorsal Striatal Neurons. J Neurosci 35:8232–8244. doi:10.1523/JNEUROSCI.1022-15.2015

Lipton DM, Gonzales BJ, Citri A. 2019. Dorsal Striatal Circuits for Habits, Compulsions and Addictions. Front Syst Neurosci 13:1–14. doi:10.3389/fnsys.2019.00028

López-López D, Gómez-Nieto R, Herrero-Turrión MJ, García-Cairasco N, Sánchez-Benito D, Ludeña MD, López DE. 2017. Overexpression of the immediate-early genes Egr1, Egr2, and Egr3 in two strains of rodents susceptible to audiogenic seizures. Epilepsy Behav 71:226–237. doi:10.1016/j.yebeh.2015.12.020

Lull ME, Freeman WM, Vrana KE, Mash DC. 2008. Correlating Human and Animal Studies of Cocaine Abuse and Gene Expression. Ann N Y Acad Sci 1141:58–75. doi:10.1196/annals.1441.013

Lüscher C. 2016. The Emergence of a Circuit Model for Addiction. Annu Rev Neurosci 39:257–276. doi:10.1146/annurev-neuro-070815-013920

Lüscher C, Malenka RC. 2011. Drug-Evoked Synaptic Plasticity in Addiction: From Molecular Changes to Circuit Remodeling. Neuron 69:650–663. doi:10.1016/j.neuron.2011.01.017

Mataga N, Fujishima S, Condie BG, Hensch TK. 2001. Experience-Dependent Plasticity of Mouse Visual Cortex in the Absence of the Neuronal Activity-Dependent Marker egr1/zif268. J Neurosci 21:9724–9732. doi:10.1523/JNEUROSCI.21-24-09724.2001

McClung CA, Nestler EJ. 2008. Neuroplasticity Mediated by Altered Gene Expression. Neuropsychopharmacology 33:3–17. doi:10.1038/sj.npp.1301544

Minatohara K, Akiyoshi M, Okuno H. 2016. Role of Immediate-Early Genes in Synaptic Plasticity and Neuronal Ensembles Underlying the Memory Trace. Front Mol Neurosci 8:78. doi:10.3389/fnmol.2015.00078

Moratalla R, Robertson H, Graybiel A. 1992. Dynamic regulation of NGFI-A (zif268, egr1) gene expression in the striatum. J Neurosci 12:2609–2622. doi:10.1523/JNEUROSCI.12-07-02609.1992

Moratalla R, Xu M, Tonegawa S, Graybiel a M. 1996. Cellular responses to psychomotor stimulant and neuroleptic drugs are abnormal in mice lacking the D1 dopamine receptor. Proc Natl Acad Sci 93:14928–14933. doi:10.1073/pnas.93.25.14928

Morita K, Okamura T, Inoue M, Komai T, Teruya S, Iwasaki Y, Sumitomo S, Shoda H, Yamamoto K, Fujio K. 2016. Egr2 and Egr3 in regulatory T cells cooperatively control systemic autoimmunity through Ltbp3-mediated TGF-β3 production. Proc Natl Acad Sci 113:E8131–E8140. doi:10.1073/pnas.1611286114

Mukherjee D, Ignatowska-Jankowska BM, Itskovits E, Gonzales BJ, Turm H, Izakson L, Haritan D, Bleistein N, Cohen C, Amit I, Shay T, Grueter B, Zaslaver A, Citri A. 2018. Salient experiences are represented by unique transcriptional signatures in the mouse brain. Elife 7:e31220. doi:10.7554/eLife.31220

Murray RC, Logan MC, Horner KA. 2015. Striatal patch compartment lesions reduce stereotypy following repeated cocaine administration. Brain Res 1618:286–298. doi:10.1016/j.brainres.2015.06.012

Nagarajan R, Svaren J, Le N, Araki T, Watson M, Milbrandt J. 2001. EGR2 Mutations in Inherited Neuropathies Dominant-Negatively Inhibit Myelin Gene Expression. Neuron 30:355–368. doi:10.1016/S0896-6273(01)00282-3

Narayana PA, Herrera JJ, Bockhorst KH, Esparza-Coss E, Xia Y, Steinberg JL, Moeller FG. 2014. Chronic cocaine administration causes extensive white matter damage in brain: Diffusion tensor imaging and immunohistochemistry studies. Psychiatry Res Neuroimaging 221:220–230. doi:10.1016/j.pscychresns.2014.01.005

Nestler E. 2002. Common Molecular and Cellular Substrates of Addiction and Memory. Neurobiol Learn Mem 78:637–647. doi:10.1006/nlme.2002.4084

Nestler EJ. 2014. Epigenetic mechanisms of drug addiction. Neuropharmacology 76:259–268. doi:10.1016/j.neuropharm.2013.04.004

Nestler EJ. 2013. Cellular basis of memory for addiction. Dialogues Clin Neurosci 15:431–43.

Nestler EJ. 2001. Molecular basis of long-term plasticity underlying addiction. Nat Rev Neurosci 2:119–128. doi:10.1038/35053570

Nestler EJ, Aghajanian GK. 1997. Molecular and Cellular Basis of Addiction. Science (80-) 278:58–63. doi:10.1126/science.278.5335.58

Nestler EJ, Hope BT, Widnell KL. 1993. Drug addiction: A model for the molecular basis of neural plasticity. Neuron 11:995–1006. doi:10.1016/0896-6273(93)90213-B

Nestler EJ, Lüscher C. 2019. The Molecular Basis of Drug Addiction: Linking Epigenetic to Synaptic and Circuit Mechanisms. Neuron 102:48–59. doi:10.1016/j.neuron.2019.01.016

Nonomura S, Nishizawa K, Sakai Y, Kawaguchi Y, Kato S, Uchigashima M, Watanabe M, Yamanaka K, Enomoto K, Chiken S, Sano H, Soma S, Yoshida J, Samejima K, Ogawa M, Kobayashi K, Nambu A, Isomura Y, Kimura M. 2018. Monitoring and Updating of Action Selection for Goal-Directed Behavior through the Striatal Direct and Indirect Pathways. Neuron 99:1302-1314.e5. doi:10.1016/j.neuron.2018.08.002

Okamura T, Sumitomo S, Morita K, Iwasaki Y, Inoue M, Nakachi S, Komai T, Shoda H, Miyazaki J, Fujio K, Yamamoto K. 2015. TGF-β3-expressing CD4+CD25−LAG3+ regulatory T cells control humoral immune responses. Nat Commun 6:6329. doi:10.1038/ncomms7329

Phillips PEM, Stuber GD, Heien MLA V., Wightman RM, Carelli RM. 2003. Subsecond dopamine release promotes cocaine seeking. Nature 422:614–618. doi:10.1038/nature01476

Piechota M, Korostynski M, Solecki W, Gieryk A, Slezak M, Bilecki W, Ziolkowska B, Kostrzewa E, Cymerman I, Swiech L, Jaworski J, Przewlocki R. 2010. The dissection of transcriptional modules regulated by various drugs of abuse in the mouse striatum. Genome Biol 11:R48. doi:10.1186/gb-2010-11-5-r48

Rakhade SN, Shah AK, Agarwal R, Yao B, Asano E, Loeb JA. 2007. Activity-dependent Gene Expression Correlates with Interictal Spiking in Human Neocortical Epilepsy. Epilepsia 48:86–95. doi:10.1111/j.1528-1167.2007.01294.x

Rebec G V., White IM, Puotz JK. 1997. Responses of neurons in dorsal striatum during amphetamine-induced focused stereotypy. Psychopharmacology (Berl) 130:343–351. doi:10.1007/s002130050249

Ribeiro EA, Scarpa JR, Garamszegi SP, Kasarskis A, Mash DC, Nestler EJ. 2017. Gene Network Dysregulation in Dorsolateral Prefrontal Cortex Neurons of Humans with Cocaine Use Disorder. Sci Rep 7:5412. doi:10.1038/s41598-017-05720-3

Rittschof CC, Hughes KA. 2018. Advancing behavioural genomics by considering timescale. Nat Commun 9:489. doi:10.1038/s41467-018-02971-0

Robison AJ, Nestler EJ. 2011. Transcriptional and epigenetic mechanisms of addiction. Nat Rev Neurosci 12:623–637. doi:10.1038/nrn3111

Rothwell PE, Fuccillo M V., Maxeiner S, Hayton SJ, Gokce O, Lim BK, Fowler SC, Malenka RC, Südhof TC. 2014. Autism-Associated Neuroligin-3 Mutations Commonly Impair Striatal Circuits to Boost Repetitive Behaviors. Cell 158:198–212. doi:10.1016/j.cell.2014.04.045

Rubio FJ, Liu Q-R, Li X, Cruz FC, Leao RM, Warren BL, Kambhampati S, Babin KR, McPherson KB, Cimbro R, Bossert JM, Shaham Y, Hope BT. 2015. Context-Induced Reinstatement of Methamphetamine Seeking Is Associated with Unique Molecular Alterations in Fos-Expressing Dorsolateral Striatum Neurons. J Neurosci 35:5625–5639. doi:10.1523/JNEUROSCI.4997-14.2015

Russo SJ, Nestler EJ. 2013. The brain reward circuitry in mood disorders. Nat Rev Neurosci 14:609–625. doi:10.1038/nrn3381

Saint-Preux F, Bores LR, Tulloch I, Ladenheim B, Kim R, Thanos PK, Volkow ND, Cadet JL. 2013. Chronic co-administration of nicotine and methamphetamine causes differential expression of immediate early genes in the dorsal striatum and nucleus accumbens of Rats. Neuroscience 243:89–96. doi:10.1016/j.neuroscience.2013.03.052

Sakaguchi M, Hayashi Y. 2012. Catching the engram: strategies to examine the memory trace. Mol Brain 5:32. doi:10.1186/1756-6606-5-32

Salery M, Trifilieff P, Caboche J, Vanhoutte P. 2020. From Signaling Molecules to Circuits and Behaviors: Cell-Type–Specific Adaptations to Psychostimulant Exposure in the Striatum. Biol Psychiatry 87:944–953. doi:10.1016/j.biopsych.2019.11.001

Savell KE, Tuscher JJ, Zipperly ME, Duke CG, Phillips RA, Bauman AJ, Thukral S, Sultan FA, Goska NA, Ianov L, Day JJ. 2020. A dopamine-induced gene expression signature regulates neuronal function and cocaine response. Sci Adv 6:eaba4221. doi:10.1126/sciadv.aba4221

Schlussman SD, Zhang Y, Kane S, Stewart CL, Ho A, Kreek MJ. 2003. Locomotion, stereotypy, and dopamine D1 receptors after chronic “binge” cocaine in C57BL/6J and 129/J mice. Pharmacol Biochem Behav 75:123–131. doi:10.1016/S0091-3057(03)00067-4

Shaham Y, Shalev U, Lu L, de Wit H, Stewart J. 2003. The reinstatement model of drug relapse: history, methodology and major findings. Psychopharmacology (Berl) 168:3–20. doi:10.1007/s00213-002-1224-x

Sinha S, Jones BM, Traniello IM, Bukhari SA, Halfon MS, Hofmann HA, Huang S, Katz PS, Keagy J, Lynch VJ, Sokolowski MB, Stubbs LJ, Tabe-Bordbar S, Wolfner MF, Robinson GE. 2020. Behavior-related gene regulatory networks: A new level of organization in the brain. Proc Natl Acad Sci 117:23270–23279. doi:10.1073/pnas.1921625117

Steiner H. 2016. Psychostimulant-Induced Gene Regulation in Striatal Circuits.Handbook of Behavioral Neuroscience. Elsevier B.V. pp. 639–672. doi:10.1016/B978-0-12-802206-1.00031-3

Steiner H, Gerfen C. 1993. Cocaine-induced c-fos messenger RNA is inversely related to dynorphin expression in striatum. J Neurosci 13:5066–5081. doi:10.1523/JNEUROSCI.13-12-05066.1993

Steiner H, Van Waes V. 2013. Addiction-related gene regulation: Risks of exposure to cognitive enhancers vs. other psychostimulants. Prog Neurobiol 100:60–80. doi:10.1016/j.pneurobio.2012.10.001

Svaren J, Meijer D. 2008. The molecular machinery of myelin gene transcription in Schwann cells. Glia 56:1541–1551. doi:10.1002/glia.20767

Terem A, Gonzales BJ, Peretz-Rivlin N, Ashwal-Fluss R, Bleistein N, del Mar Reus-Garcia M, Mukherjee D, Groysman M, Citri A. 2020. Claustral Neurons Projecting to Frontal Cortex Mediate Contextual Association of Reward. Curr Biol 30:3522-3532.e6. doi:10.1016/j.cub.2020.06.064

Tonegawa S, Pignatelli M, Roy DS, Ryan TJ. 2015. Memory engram storage and retrieval. Curr Opin Neurobiol 35:101–109. doi:10.1016/j.conb.2015.07.009

Topilko P, Schneider-Maunoury S, Levi G, Baron-Van Evercooren A, Chennoufi ABY, Seitanidou T, Babinet C, Charnay P. 1994. Krox-20 controls myelination in the peripheral nervous system. Nature 371:796–799. doi:10.1038/371796a0

Turm H, Mukherjee D, Haritan D, Tahor M, Citri A. 2014. Comprehensive Analysis of Transcription Dynamics from Brain Samples Following Behavioral Experience. J Vis Exp 1–12. doi:10.3791/51642

Tyssowski KM, DeStefino NR, Cho J-H, Dunn CJ, Poston RG, Carty CE, Jones RD, Chang SM, Romeo P, Wurzelmann MK, Ward JM, Andermann ML, Saha RN, Dudek SM, Gray JM. 2018. Different Neuronal Activity Patterns Induce Different Gene Expression Programs. Neuron 98:530-546.e11. doi:10.1016/j.neuron.2018.04.001

Tyssowski KM, Gray JM. 2019. The neuronal stimulation–transcription coupling map. Curr Opin Neurobiol 59:87–94. doi:10.1016/j.conb.2019.05.001

Valjent E. 2006. Plasticity-Associated Gene Krox24/Zif268 Is Required for Long-Lasting Behavioral Effects of Cocaine. J Neurosci 26:4956–4960. doi:10.1523/JNEUROSCI.4601-05.2006

Voiculescu O, Charnay P, Schneider-Maunoury S. 2000. Expression pattern of aKrox-20/Cre knock-in allele in the developing hindbrain, bones, and peripheral nervous system. genesis 26:123–126. doi:10.1002/(SICI)1526-968X(200002)26:2<123::AID-GENE7>3.0.CO;2-O

Volkow ND, Morales M. 2015. The Brain on Drugs: From Reward to Addiction. Cell 162:712–725. doi:10.1016/j.cell.2015.07.046

Volkow ND, Wang G-J, Telang F, Fowler JS, Logan J, Childress A-R, Jayne M, Ma Y, Wong C. 2006. Cocaine Cues and Dopamine in Dorsal Striatum: Mechanism of Craving in Cocaine Addiction. J Neurosci 26:6583–6588. doi:10.1523/JNEUROSCI.1544-06.2006

Walker DM, Cates HM, Loh Y-HE, Purushothaman I, Ramakrishnan A, Cahill KM, Lardner CK, Godino A, Kronman HG, Rabkin J, Lorsch ZS, Mews P, Doyle MA, Feng J, Labonté B, Koo JW, Bagot RC, Logan RW, Seney ML, Calipari ES, Shen L, Nestler EJ. 2018. Cocaine Self-administration Alters Transcriptome-wide Responses in the Brain’s Reward Circuitry. Biol Psychiatry 84:867–880. doi:10.1016/j.biopsych.2018.04.009

Warner LE. 1999. Functional consequences of mutations in the early growth response 2 gene (EGR2) correlate with severity of human myelinopathies. Hum Mol Genet 8:1245–1251. doi:10.1093/hmg/8.7.1245

Warner LE, Mancias P, Butler IJ, McDonald CM, Keppen L, Koob KG, Lupski JR. 1998. Mutations in the early growth response 2 (EGR2) gene are associated with hereditary myelinopathies. Nat Genet 18:382–384. doi:10.1038/ng0498-382

White IM, Doubles L, Rebec G V. 1998. Cocaine-induced activation of striatal neurons during focused stereotypy in rats. Brain Res 810:146–152. doi:10.1016/S0006-8993(98)00905-6

Wilkinson DG. 1995. Genetic control of segmentation in the vertebrate hindbrain. Perspect Dev Neurobiol 3:29–38.

Wolf ME. 2016. Synaptic mechanisms underlying persistent cocaine craving. Nat Rev Neurosci 17:351–365. doi:10.1038/nrn.2016.39

Worley P, Bhat R, Baraban J, Erickson C, McNaughton B, Barnes C. 1993. Thresholds for synaptic activation of transcription factors in hippocampus: correlation with long-term enhancement. J Neurosci 13:4776–4786. doi:10.1523/JNEUROSCI.13-11-04776.1993

Yamada K, Gerber DJ, Iwayama Y, Ohnishi T, Ohba H, Toyota T, Aruga J, Minabe Y, Tonegawa S, Yoshikawa T. 2007. Genetic analysis of the calcineurin pathway identifies members of the EGR gene family, specifically EGR3, as potential susceptibility candidates in schizophrenia. Proc Natl Acad Sci 104:2815–2820. doi:10.1073/pnas.0610765104

Yap E-L, Greenberg ME. 2018. Activity-Regulated Transcription: Bridging the Gap between Neural Activity and Behavior. Neuron 100:330–348. doi:10.1016/j.neuron.2018.10.013

Yin HH, Knowlton BJ, Balleine BW. 2004. Lesions of dorsolateral striatum preserve outcome expectancy but disrupt habit formation in instrumental learning. Eur J Neurosci 19:181–189. doi:10.1111/j.1460-9568.2004.03095.x

Zahm DS, Becker ML, Freiman AJ, Strauch S, DeGarmo B, Geisler S, Meredith GE, Marinelli M. 2010. Fos After Single and Repeated Self-Administration of Cocaine and Saline in the Rat: Emphasis on the Basal Forebrain and Recalibration of Expression. Neuropsychopharmacology 35:445–463. doi:10.1038/npp.2009.149

Zapata A, Minney VL, Shippenberg TS. 2010. Shift from Goal-Directed to Habitual Cocaine Seeking after Prolonged Experience in Rats. J Neurosci 30:15457–15463. doi:10.1523/JNEUROSCI.4072-10.2010

